# Multivalency, autoinhibition, and protein disorder in the regulation of interactions of dynein intermediate chain with dynactin and the nuclear distribution protein

**DOI:** 10.1101/2022.06.05.494868

**Authors:** Kayla A. Jara, Nikolaus M. Loening, Patrick N. Reardon, Zhen Yu, Prajna Woonnimani, Coban Brooks, Cat H. Vesely, Elisar J. Barbar

## Abstract

Cytoplasmic dynein plays crucial roles in the intracellular transport of organelles and other cargoes. Central to dynein function is the intrinsically disordered N-terminal domain of dynein intermediate chain (IC), which binds the three dimeric dynein light chains at multivalent sites, and dynactin p150^Glued^ and nuclear distribution protein (NudE) at overlapping sites. The disorder in IC has hindered cryo-electron microscopy and X-ray crystallography studies of its structure and interactions. Here we use a suite of biophysical methods to reveal how multivalent binding of the three light chains regulate IC interactions with p150^Glued^ and NudE. Using the N-terminal domain or the full-length IC from *Chaetomium thermophilum*, a tractable species to interrogate IC interactions, we identify a significant reduction in IC’s binding affinity for p150^Glued^ and a loss of binding to NudE in contrast to the tight binding observed with small IC constructs. We attribute this difference to autoinhibition caused by strong long-range intramolecular interactions that cover IC’s N-terminal single α-helix, the site for p150^Glued^ and NudE binding. Reconstitution of IC subcomplexes demonstrate that autoinhibition is differentially regulated by light chains binding underscoring their importance both in assembly and organization of IC, and in selection between multiple binding partners at the same site.

## Introduction

Dynein intermediate chain (IC) plays key roles in modulating dynein interactions and activity (1–5). For example, IC connects the three dynein light chains (Tctex, LC8, and LC7) to the heavy chain, and also serves as the primary binding site of multiple non-dynein proteins essential for dynein regulation, such as the p150^Glued^ subunit of dynactin. Despite this level of importance, high resolution structural information of IC interactions is limited due to its highly disordered N-terminal domain, which hinders studies by methods such as X-ray crystallography and cryo-electron microscopy (cryo-EM).

The primarily disordered N-terminus of IC (N-IC) (4) contains a single α-helix (SAH) and a short helix (H2), each of which is followed by disordered linker regions (3), whereas the C-terminus of IC (C-IC) folds into a β-propeller and contains the binding site for the dynein heavy chain (6, 7) (Fig 1A). Apo-IC is monomeric but, upon binding the homodimeric dynein light chain subunits (Tctex, LC8, and LC7), it dimerizes to form a subcomplex that is best described as a polybivalent scaffold (5,8–12). In the formation of this subcomplex, the first binding event pays the entropic cost for subsequent bivalent binding events (8,13,14) and the enhancement of subsequent binding events is modulated by the length of the disordered linkers separating the binding sites (14, 15). Such a mechanism has been well-described for the IC/Tctex/LC8 complex, for which a three-residue linker separates the Tctex and LC8 binding sites. In this complex, binding of one light chain to IC results in a 50-fold enhancement in the affinity for binding the other light chain (14). Each homodimeric light chain has a corresponding binding site on IC that is initially disordered but forms β-strands (for Tctex and LC8) or an α-helix (for LC7) when bound and incorporated into the fold of their respective ligand. The assembly of monomeric IC and the homodimeric light chains is such that, when bound, folding occurs only at the protein-protein interfaces (12, 14) while the remaining linker regions stay completely disordered (13,16,17). In this work, we demonstrate the importance of the flexibility of these disordered linker regions separating the three dimeric light chains in regulating interactions of IC with non-dynein binding partners.

**Figure 1:**
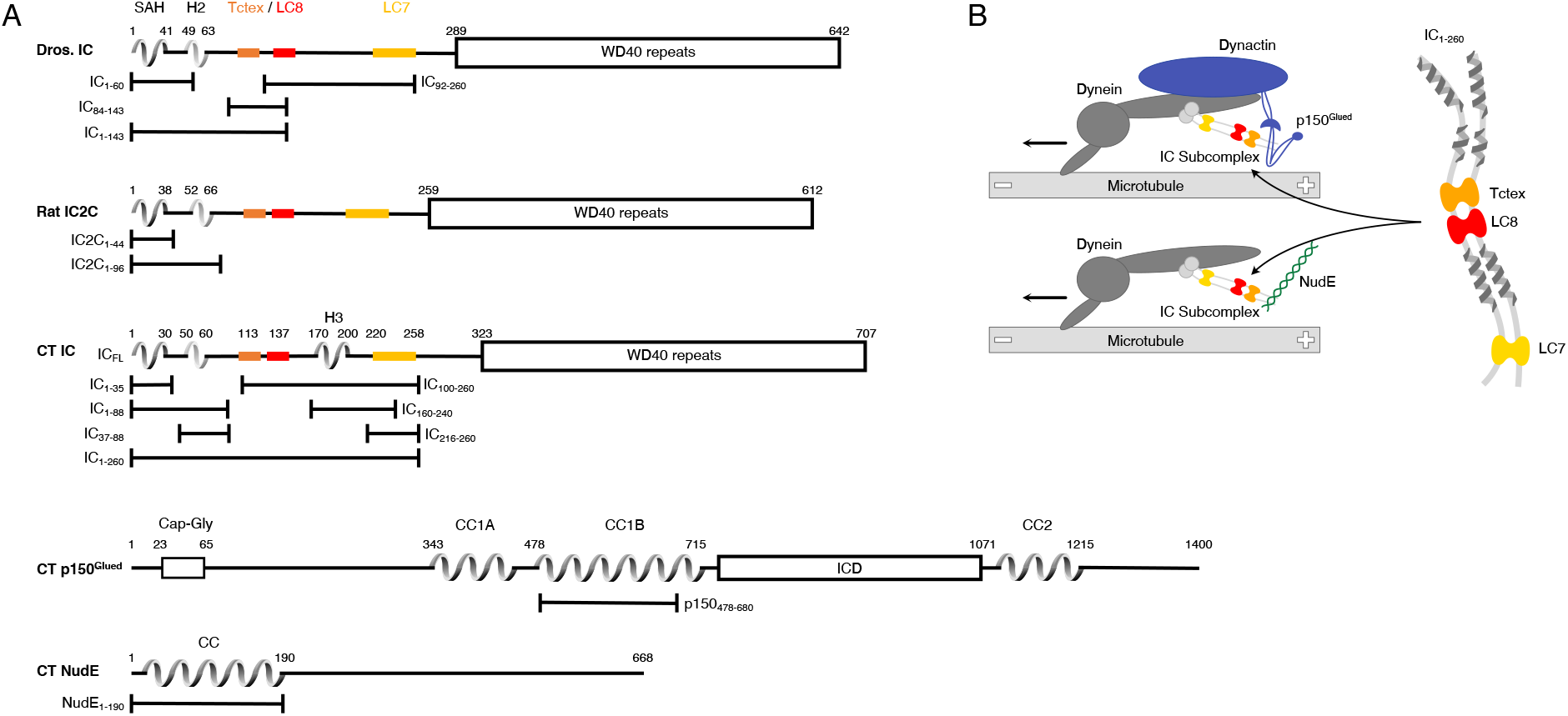
Domain architecture for dynein intermediate chain (IC), dynactin p150^Glued^, and nuclear distribution protein (NudE). (A) Domain architecture diagrams for IC from *Drosophila melanogaster* (Dros. IC) and *Rattus norvegicus* (Rat IC2C) and constructs used in earlier work are provided for comparison. Proteins and constructs used in this work are from *Chaetomium thermophilum* (CT). All IC’s have an N-terminal single α-helix (SAH), followed by either a transient/nascent or folded second helix (H2). In CT, there is an additional helix (H3). The Tctex (orange), LC8 (red), and LC7 (yellow) binding sites are well characterized in Dros. IC and Rat IC2C, and their position in CT was predicted based on sequence and structure comparison. The C-terminal domain is predicted to contain seven WD40 repeats. The CT constructs IC_FL_, IC_1-88_, IC_37-88_, IC_1-260_, IC_100-260_, IC_140-260_, IC_216-260_, and IC_216-260_ are used in this paper; the IC_1-35_ construct was used in prior work. CT p150^Glued^ is predicted to have a Cap-Gly domain near the N-terminus, and two coiled-coil domains, CC1 and CC2, that are separated by an intercoil domain (ICD). CC1 is further divided into two regions called CC1A and CC1B. p150_478-680_ (p150_CC1B_) is the construct used in this work. CT NudE is predicted to have an N-terminal coil-coiled (CC) region followed by disorder. NudE_1-190_ (NudE_CC_) is the construct used in this work. (B) Contextual models of dynein (dark grey) with the IC subcomplex highlighted (light grey). The top model depicts the interaction between N-IC and the p150^Glued^ subunit of dynactin (blue) while the bottom model depicts the interaction between N-IC and NudE (green). In both models, dynein is a processive motor traveling towards the minus end of a microtubule. For clarity, shown on a larger scale to the right is the IC subcomplex consisting of CT IC_1-260_ with the homodimeric dynein light chains: Tctex (orange), LC8 (red), and LC7 (yellow).

N-IC binds non-dynein regulatory proteins including dynactin and nuclear distribution protein (NudE) (2,3,18–22). Dynactin is a multisubunit complex that binds dynein with its largest subunit, p150^Glued^, and this interaction is required for the recruitment of cargo, dynein processivity, and correct spindle formation in cell division (19,23–28). NudE, on the other hand, regulates dynein recruitment to kinetochores and membranes, centrosome migration, mitotic spindle orientation, and binds LIS1 (24,29–35). It is well recognized that IC partnering with either p150^Glued^ or NudE impacts the regulation of the dynein complex as this dictates its interactions with adaptors and cargo (1). However, the molecular processes underlying which regulator is bound at any given time are still unclear. Previously, we showed that in multiple species (rat, Drosophila, yeast) the coiled-coil domains of p150^Glued^ and NudE from the respective species each bind IC at the same site (the SAH region) (2,20,21) but that in a filamentous fungus *(Chaetomium thermophilum)* binding of IC and p150^Glued^ also involves binding of the H2 region (22).

With advancements in cryo-EM, the overall picture of dynein structure and activity is beginning to emerge. For example, cryo-EM images show that dynactin causes the motor domains of dynein to reorient to become parallel to microtubules prior to binding (7, 36). Additionally, adaptors can recruit a second dynein to dynactin for faster movement (6). Dynein is seemingly a perfect candidate for structural characterization by cryo-EM, as the motor domains are large and symmetric. However, a recurring theme in these studies is that the flexibility of N-IC limits details of its structure and binding interactions with multiple regulators. Although residue specific studies have been performed on IC, they are thus far limited to short fragments and to conditions far removed from native biological systems. Using the combined data from EM structures and *in vitro* studies, we have made a model to aid in visualizing the assembled N-IC subcomplex bound to either p150^Glued^ or NudE thus providing context for the work presented here (Fig 1B).

The interactions of short fragments of N-IC with dynein light chains and non-dynein proteins from multiple different species (rat, Drosophila, yeast) demonstrate that NudE and p150^Glued^ compete for the same binding site, however a mechanism for IC partner selection and the importance of bivalency in IC subcomplex assembly have remained elusive (2,3,12,14,20,21,30,31,37). We recently introduced *Chaetomium thermophilum* (CT), a thermophilic filamentous fungus, as a new system for dynein studies (22). The CT IC_1-260_ (residues 1-260) construct used in this work is far longer than our previously studied CT, Drosophila, and rat constructs (Fig 1, S1). Utilizing the entire N-terminal domain of IC allows, for the first time, characterization of IC interactions in the context of its assembly with the light chains, as has been shown vital for other disorder-driven systems (38–42). However, none of the previously studied systems have the level of complexity (an assembly of five unique proteins) as the system explored here. From our studies of both IC_1-260_ as well as full-length IC, we 1) describe the first recombinant expression and reconstitution of the polybivalent scaffold formed from IC_1-260_ bound by all three light chains (Tctex, LC8, and LC7), as well as by coiled-coil domains of p150^Glued^ or NudE (Fig 1), 2) we identify long range tertiary contacts between residues in the C-terminal region (the LC7 binding site) and residues in the N-terminal region (SAH) that inhibit binding to non-dynein proteins 3) we show how assembly with the light chains relieves this autoinhibition and regulates binding of IC to p150^Glued^ and NudE and 4) we demonstrate for the first time the essential role of light chains in both assembly and regulation of the full-length IC.

## Results

### CT p150_CC1B_ and NudE_CC_ are dimeric, while CT IC_1-260_ is monomeric

Studies of interactions of IC constructs with light chains show that two monomeric IC chains are brought together by the dimeric light chains to create a ‘ladder-like’ polybivalent scaffold (8,13,14,17). Here we use a construct of IC that includes all the binding sites for the light chains and non-dynein proteins, the full-length light chains, and coiled-coil domains of non-dynein proteins and employ multiple techniques to determine their association states. Sedimentation velocity analytical ultracentrifugation (SV-AUC), which gives information about a protein’s mass and shape in solution, was used to determine each protein’s heterogeneity and size. A larger sedimentation coefficient (S) indicates a protein with a larger mass and/or a protein with a smaller, shape dependent, frictional ratio (43). SV-AUC reveals that IC_1-260_, the three light chains, and non-dynein proteins p150_CC1B_ and NudE_CC_ all have similar sedimentation coefficients (in the 2-3 S range, Fig 2A). Further, each subunit shows a single sharp peak in the c(S) distribution, indicating that the proteins are homogeneous in solution. In comparison to the sedimentation coefficients for IC_1-260_ and the light chains, both p150_CC1B_ and NudE_CC_ have smaller sedimentation coefficients than would typically be expected for globular proteins with their respective dimeric masses. However, as sedimentation coefficients also depend on shape, the smaller values observed for p150_CC1B_ and _CC_ are consistent with their predicted coiled-coil structures (which result in elongated, less-compact, rod-like shapes) causing slower sedimentation than if they were globular.

**Figure 2:**
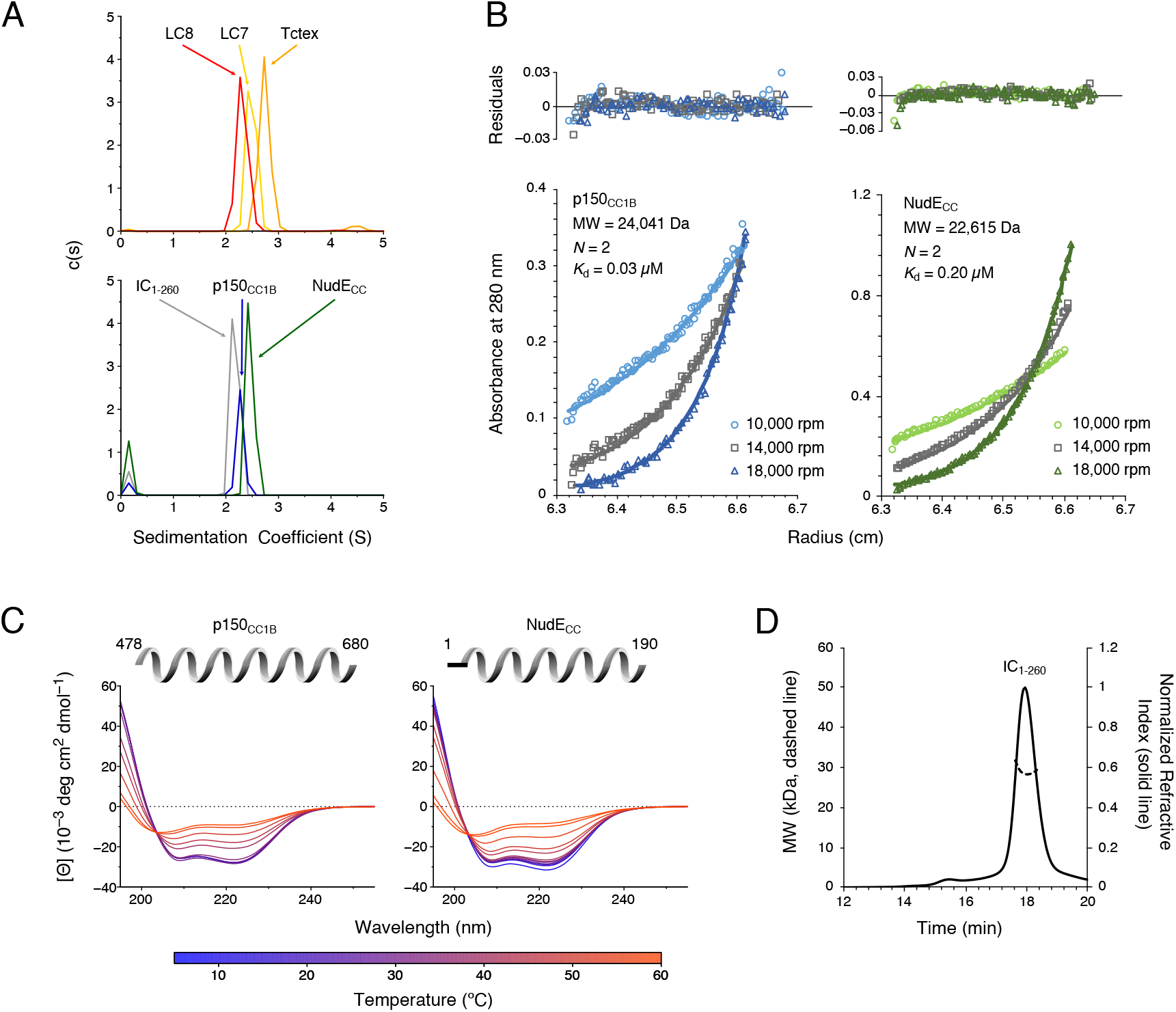
CT p150_CC1B_, NudE_CC_, and dynein light chains are dimeric, whereas CT IC_1-260_ is monomeric. (A) SV-AUC profiles for LC8 (red), LC7 (yellow), and Tctex (orange) (top), and, IC_1-260_ (grey), p150_CC1B_ (blue), and NudE_CC_ (green) (bottom). All samples were at protein concentration of 30 µM. (B) SE-AUC data for p150_CC1B_ (blue) and NudE_CC_ (green) at three speeds (10,000, 14,000, and 18,000 rpm). Data were fit to a monomer-dimer binding model. The quality of the fits to this model is reflected by the plots of the residuals on top. The monomeric masses determined by fitting this data compare very well to the masses expected based on the sequences for the constructs. The stoichiometry (*N*) values of 2 indicate that both p150_CC1B_ and NudE_CC_ are dimers in solution. (C) CD spectra of p150_CC1B_ and NudE_CC_ acquired at temperatures in the 5 to 60°C range. The shape of the spectra for both p150_CC1B_ and NudE_CC_ indicate α-helical secondary structure, and the 222/208 ratios (1.04 and 1.00 for p150_CC1B_ and NudE_CC_, respectively) are consistent with coil-coiled structures. (D) MALS of IC_1-260_ gives an estimated mass of 29.5 kDa, which indicates that on its own, IC_1-260_ exists as a monomer in solution (calculated mass of monomer is 29.2 kDa). **Figure 2-source data 1: Source files for SV-AUC, SE-AUC, CD, and SEC-MALS data.** This Excel workbook contains all the data plotted in Figure 2. The different sheets correspond to different panels within the figure. Additional information regarding data collection can be found in the corresponding methods sections. Data were plotted using gnuplot.

Sedimentation equilibrium analytical ultracentrifugation (SE-AUC) confirmed the dimeric coiled-coil state of p150_CC1B_ and NudE_CC_. SE-AUC data fit to a monomer-dimer binding model results in dimerization dissociation constants of 0.03 μM and 0.20 μM for p150_CC1B_ and NudE_CC_, respectively, indicating that both are strong dimers and that, of the two, p150_CC1B_ is the tighter dimer (Fig 2B). This result is underscored by circular dichroism (CD) spectra acquired at temperatures ranging from 5-60°C (Fig 2C), which show that while both proteins have α-helical secondary structure (44), they have different mechanisms of unfolding. Changes in the secondary structure for p150_CC1B_ are most dramatic between 30 and 35°C, whereas NudE_CC_ shows gradual unfolding across the 5-40°C temperature range. Also of note, the coiled-coil structures of p150_CC1B_ and NudE_CC_ are confirmed by their CD spectra which show helical structures and values for 222/208 ratios larger than 1 (Fig 2C).

To confirm our expectation that IC_1-260_ is a monomer in solution, we used size exclusion chromatography (SEC) with multi-angle light scattering (MALS) detection (Fig 2D). The measured mass of approximately 29.5 kDa based on the MALS data is consistent with the expected mass of 29.2 kDa for an IC_1-260_ monomer.

### CT IC_1-260_ is stabilized by long range contacts

The N-IC from CT has a domain architecture with structural elements that are similar in all experimentally characterized ICs: an N-terminal single α-helix (SAH), a nascent helix 2 (H2), and long disordered linkers (Fig 1A). CT IC is unique, however, in also including a strongly predicted third helix (H3) corresponding to residues 170-200 (45–51) (Fig 3A, S2). To validate this predicted secondary structure, and determine its impact on global stability, we acquired CD spectra of various IC constructs: IC_1-260_, IC_1-88_ (containing SAH and H2), IC_100-260_ (containing linker, H3, and the LC7 binding site), and IC_160-240_ (containing H3 and shorter linker). All the CD spectra show two minima around 208 and 222 nm, which is indicative of the presence of α-helical secondary structure (44) (Fig 3B-E). The estimated fractional helicity (52) values for these constructs at 5°C is in the range of 20%-35%, which matches the fraction of either NMR-determined or predicted helical residues in each construct (Fig 3A). Notably, the longest IC construct (IC_1-260_) is the most thermostable of the four, and resists unfolding until above 50°C. Comparatively, IC_1-88_, IC_160-240_, and IC_100-260_ exhibit some loss in secondary structure significantly below 50°C. In particular, IC_100-260_ appears to be the least stable of the three, with a gradual loss of secondary structure that begins around 25°C and a complete loss of helical structure around 40°C. From this, it is reasonable to hypothesize that some degree of tertiary contacts may exist only within IC_1-260_ and underlie the increase in its structural stability and cooperative unfolding.

**Figure 3:**
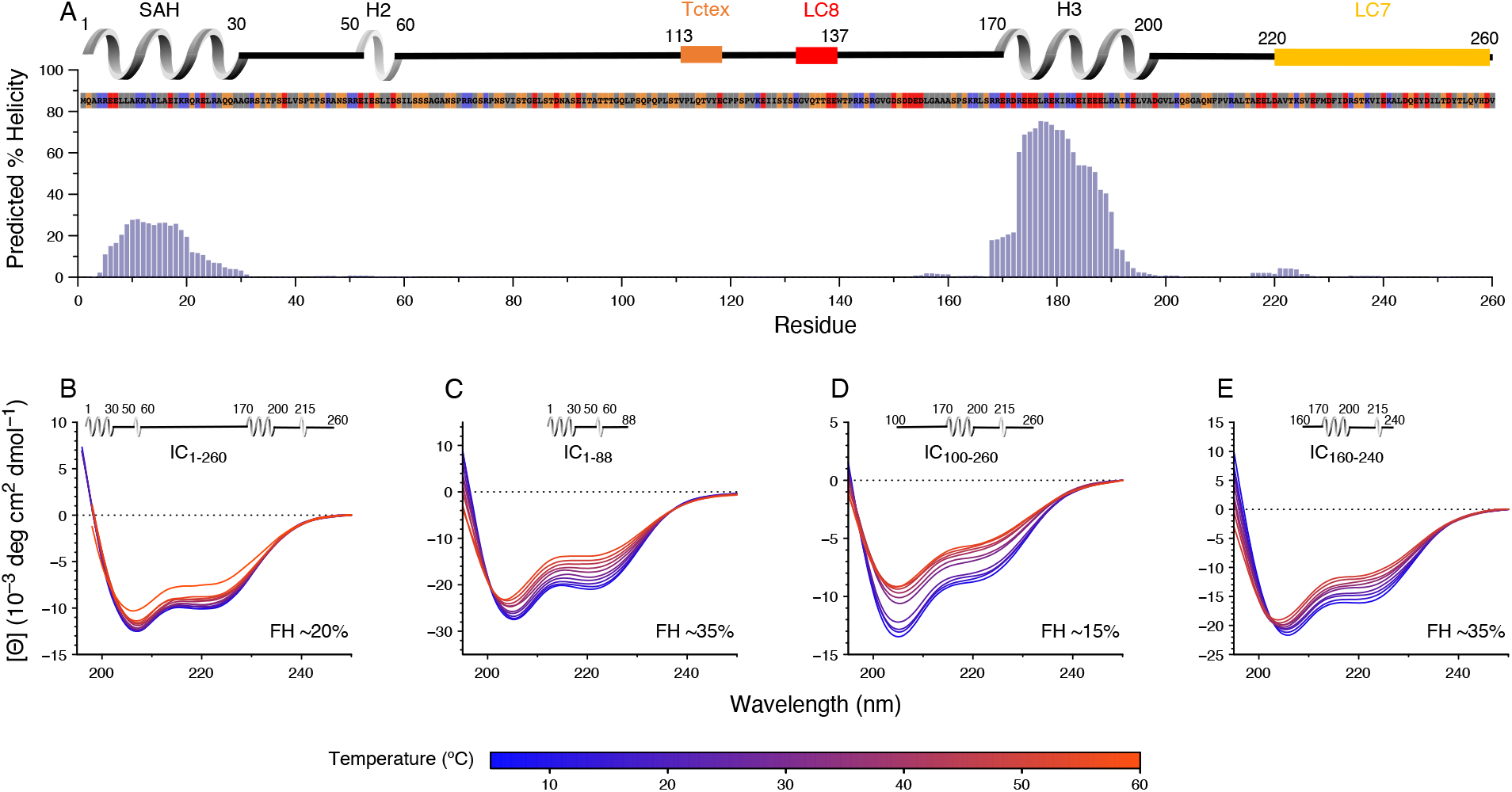
Secondary structure and thermal stability of CT IC. (A) Agadir prediction for IC_1-260_ showing the percent helicity by residue (purple). Shown above the plot is a schematic structure for IC_1-260_ with labels for SAH, H2 and H3 above the helical structure. The sites for lights chains binding are also indicated. The amino acid sequence under the schematic is colored by amino acid type: hydrophobic (grey), positive (red), negative (blue), neutral (orange). Variable temperature CD spectra of (B) IC_1-260_, (C) IC_1-88_, (D) IC_100-260_, and (E) IC_160-240_. The shapes of the spectra for all constructs indicates a mixture of α-helical secondary structure and regions of intrinsic disorder. Loss in structure, or lack thereof, over a temperature range of 5-50°C (blue for lowest, red for highest) indicates how each construct varies in stability and indicates that IC_1-260_ is the most thermally stable. The fractional helicity (FH) of each construct at 5°C was calculated based on the experimentally observed mean residue ellipticity at 222 nm as explained in the methods section. **Figure 3-source data 2: Source files for IC_1-260_ helicity prediction and CD data.** This Excel workbook contains all the data plotted in Figure 3. The different sheets correspond to different panels within the figure. Additional information regarding data collection can be found in the corresponding methods sections. Data were plotted using gnuplot.

To further probe possible tertiary contacts within IC_1-260_, SV-AUC was employed to determine if there is binding between the IC_1-88_ and IC_100-260_ constructs (Fig S3 A). IC_1-260_ has a sedimentation coefficient of approximately 2.2 S, whereas IC_100-260_ has a smaller sedimentation coefficient of 1.5 S due to its lower mass and its expected elongation compared to IC_1-260_ (Fig S3 B). Upon addition of IC_1-88_ at a 1:2 molar ratio (IC_100-260_:IC_1-88_), a complex with a sedimentation coefficient of 2 S is formed, indicating a strong interaction between the two constructs; a peak for the excess of IC_88_ is not observed because IC_1-88_ does not absorb at 280 nm. The IC_1-88_/IC_100-260_ complex has a slightly smaller sedimentation coefficient than that of IC_1-260_, which can be explained by a greater degree of elongation for this complex compared to IC_1-260_. SEC-MALS determined a mass of 30.3 kDa for the IC_1-88_/IC_100-260_ complex which matches the expected mass of 30.6 kDa for a 1:1 complex (Fig S3 C). Together, these data confirm the presence of tertiary contacts within IC_1-260_ and explain the increase in its stability compared to smaller IC constructs (Fig 3).

### Identifying disordered domains of CT IC in the context of IC_1-260_

To identify the disordered regions of IC_1-260_, we used nuclear magnetic resonance (NMR) spectroscopy to study isotopically-labeled protein samples. The limited chemical shift dispersion in the ^1^H-^15^N TROSY spectrum for IC_1-260_ at 10°C (Fig 4B), along with appearance of the majority of the peaks in the CLEANEX experiment at this temperature (Fig 4C), indicate that the peaks observed in the spectra are for the disordered regions of the protein. Using triple resonance experiments on a ^2^H/^13^C/^15^N labeled sample, we assigned almost all of the observable peaks, corresponding to 37% (90 of 245) of the non-proline residues (Fig 4B). The handful of “unassigned” peaks in Fig 5B mainly correspond to side-chain amides or to peaks from minor conformers that arise due to the slow cis/trans isomerization of the peptide bond between prolines and the amino acid preceding them (53). The assigned peaks all correspond to residues in disordered linker regions of IC_1-260_ and these peaks vary considerably in intensity (Fig 4A).

**Figure 4:**
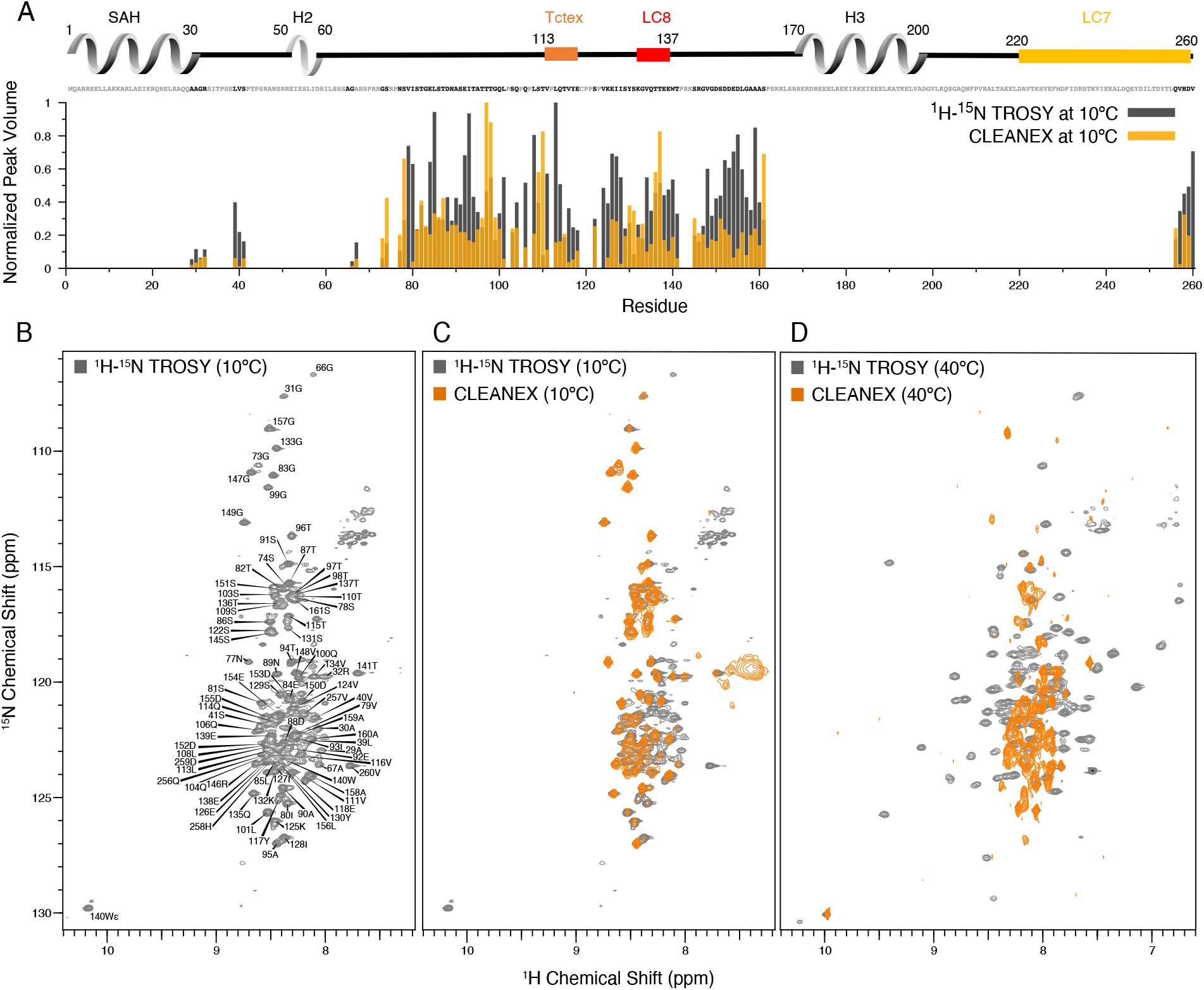
Identification of disordered linkers of CT IC_1-260_ using NMR spectroscopy. (A) Plot showing the normalized peak volumes at 10°C in the ^1^H-^15^N TROSY spectrum (grey) and in the CLEANEX spectrum (orange) of the amides that could be assigned. Assigned residues are in black in the sequence above the plot (and unassigned residues in grey); all assigned residues are from disordered regions of IC_1-260_. (B) ^1^H-^15^N TROSY spectrum of IC_1-260_ acquired at 800 MHz at 10°C showing amide assignments. (C) Overlay of a CLEANEX spectrum (orange) with the ^1^H-^15^N TROSY spectrum (grey) at 10°C, shows that most of the assignable residues are in exchange with the solvent on the timescale of the CLEANEX experiment. (D) At 40°C, ^1^H-^15^N TROSY spectrum (grey) shows new peaks appearing with greater chemical shift dispersion for IC_1-260_ in the 800 MHz. Overlaying a CLEANEX spectrum (orange) at this temperature reveals that most of the new peaks in the ^1^H-^15^N TROSY spectrum are from amides that are slow to exchange with the solvent and therefore are not observed in the CLEANEX spectrum. **Figure 4-source data 3: Source files for IC_1-260_ NMR TROSY and CLEANXEX data.** This Excel workbook contains the peak heights and volumes for residues within IC_1-260_ in the TROSY and CLEANEX NMR experiments collected at 10°C. Additional information regarding data collection can be found in the corresponding methods section. Data were plotted using gnuplot.

**Figure 5:**
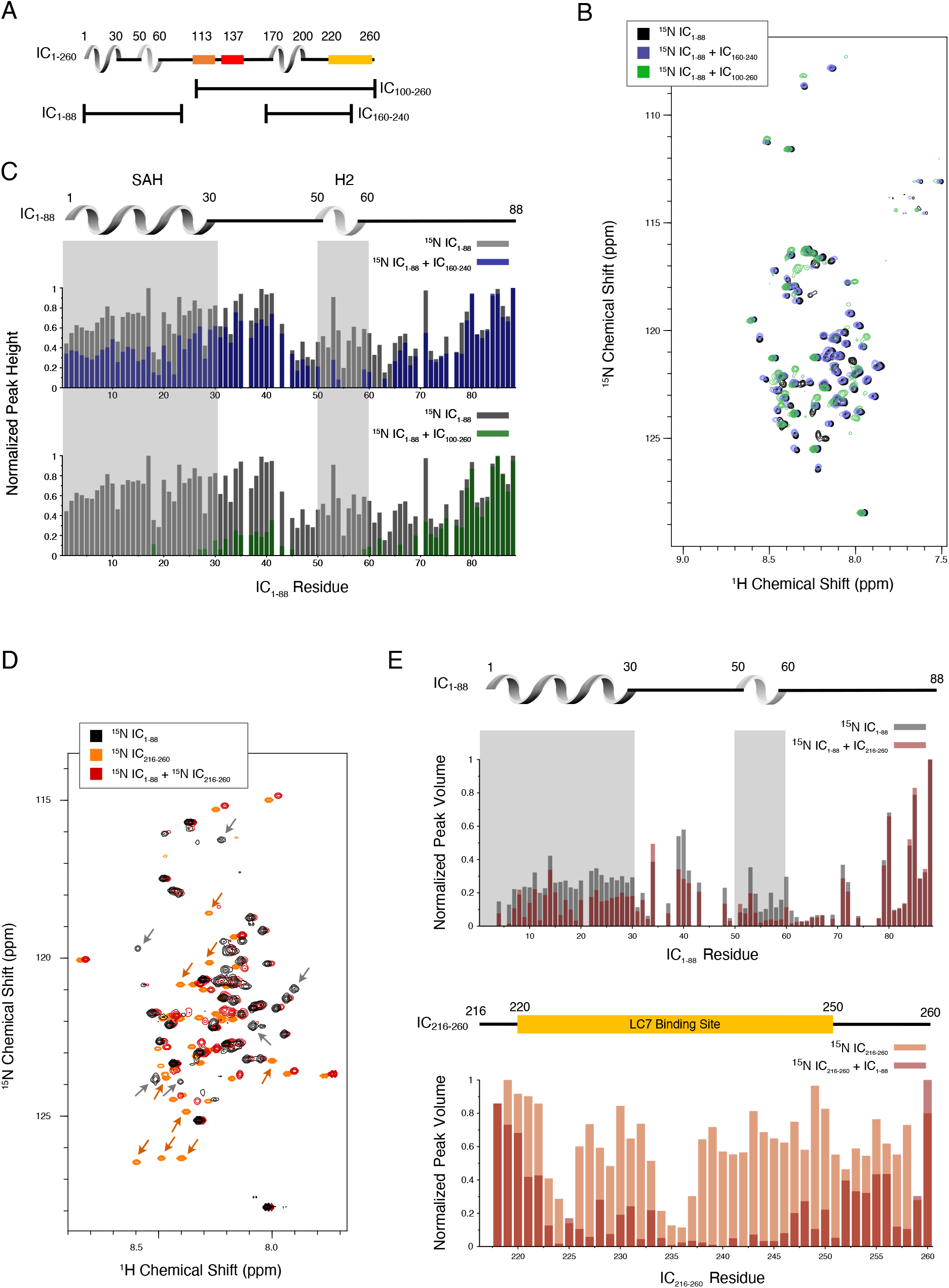
Evidence of tertiary contacts between the N and C-termini within CT IC_1-260_. (A) Domain architecture diagram for CT IC_1-260_ with bars shown below corresponding to the IC_100-260_, IC_1-88_, and IC_160-240_ constructs. (B) ^1^H-^15^N TROSY overlays of free ^15^N-labeled IC_1-88_ (black) and ^15^N-labeled IC_1-260_ bound to unlabeled IC_160-240_ (purple) and IC_100-260_ (green). Note, spectra are deliberately offset in the ^1^H dimension to help visualize overlapping peaks. (C) Normalized peak heights in the ^1^H-^15^N TROSY spectra for (top) ^15^N-labeled IC_1–88_ (grey) and ^15^N-labeled IC_1–88_ + IC_160-240_ (purple) and (bottom) ^15^N-labeled IC_1–88_ (grey) and ^15^N-labeled IC_1–88_ + IC_100-260_ (green). (D) ^1^H-^15^N HSQC overlay of ^15^N-labeled IC_1-88_ (black), ^15^N-labeled IC_216-260_ (orange), and ^15^N-labeled IC_216-260_ bound to ^15^N-labeled IC_1-88_ (red). Arrows highlight some of the more significant peak disappearances for IC_1-88_ (grey arrows) and IC_216-260_ (orange arrows). Note, spectra are deliberately offset by 0.03 ppm in the ^1^H dimension to help visualize overlapping peaks. (E) Normalized peak volumes in the ^1^H-^15^N HSQC spectra for ^15^N-labeled IC_216-260_ (top, grey columns) and ^15^N-labeled IC_216-260_ (bottom, orange columns) when free and when in the presence of the other protein (red columns). **Figure 5-source data 4: Source files for NMR binding studies of IC fragments.** This Excel workbook contains the peak heights and volumes for residues within the ^15^N-labeled IC_1-88_ and ^15^N-labeled IC_216-260_ constructs upon binding of IC_100-260_, IC_160-240_, and ^15^N-labeled IC_216-260_ or ^15^N-labeled IC_1-88_. Additional information regarding data collection can be found in the corresponding methods section. Data were plotted using gnuplot.

By increasing the temperature to 40°C, peaks with much greater chemical shift dispersion become visible in the ^1^H-^15^N TROSY spectrum (Fig 4D), while peaks corresponding to disordered regions of the protein disappear. Based on CD data (Fig 3B), IC_1-260_ does not undergo significant secondary structural changes in the 10 to 40°C temperature range. Therefore, the newfound peak dispersion at higher temperature is not due to an increase in secondary structure. Rather, the appearance of peaks at elevated temperatures is most likely due to an increased rate of molecular tumbling causing peaks from more structured parts of the protein that were too broad to observe at lower temperatures to narrow and become visible. The conclusion that these peaks belong to residues in ordered regions is supported by their chemical shift dispersion and their absence in the CLEANEX spectrum at 40°C (Fig 4D). Some additional peaks are observed in the CLEANEX spectrum at 40°C that do not appear in the TROSY spectrum at this temperature; these correspond to amides in disordered regions of the protein that are in exchange with the solvent with exchange rates that makes them detectable in the CLEANEX experiment, but invisible in the TROSY experiment. Together, these spectra support the conclusion that IC_1-260_ contains both structured and disordered regions, with the most disordered regions resulting in peaks that are the most easily assigned by NMR at low temperature. Even with the use of deuterated samples and TROSY-based experiments, the peaks at 40°C corresponding to more structured regions were, for the most part, not assignable as their rapid relaxation resulted in extremely low signal intensities in backbone assignment experiments. IC_1-260_ samples with salt concentrations of 20 mM and 250 mM were also explored at both 10°C and 40°C, to ensure that electrostatic interactions were not the cause for missing peaks (Fig S4).

### The compact structure of CT IC_1-260_

To identify the residues involved in the tertiary contacts within IC_1-260_, we collected NMR data on smaller fragments of IC; IC_1-88_, IC_100-260_, and IC_160-240_ (Fig 1, 5A). Based on the long-range interactions observed between IC_1-88_ and IC_100-260_ (Fig 3B, S3), we hypothesized that an intramolecular interaction between the SAH region (residues 1-30), and the H3 region (residues 170-200) could be contributing to the stability of IC_1-260_. To test this, we added unlabeled IC_160-240_ to ^15^N-labeled IC_1-88_. Peak disappearances and peak shifts for the spectrum of IC_1-88_ upon addition of IC_160-240_ indicate some degree of interaction (Fig 5B-C). However, when conducting the same experiment with unlabeled IC_100-260_ instead of unlabeled IC_160-240_, the effects were much more pronounced (Fig 5B-C). In both cases, peaks corresponding to residues in the SAH and H2 regions of IC_1-88_ either lost intensity or disappeared completely, while peaks from linker region residues remained largely unaffected, especially those that are near the C-terminus of IC_1-88_.

To continue to narrow down the location of the interaction, an additional IC construct, IC_216-260_, that does not include the H3 region, was ^15^N-labeled and completely assigned. Upon titration with ^15^N-labeled IC_1-88_, many of the IC_216-260_ peaks disappear or shift (Fig 5D-E). From this, we conclude that the tertiary contacts within IC_1-260_ are largely between the SAH/H2 regions and a region near the C-terminus (residues 220-250) that overlaps with the LC7 binding site. We note that these interactions appear to be even stronger in the context of the longer disordered chain encompassing the Tctex and LC8 binding sites based on the more substantial disappearances of peaks from the SAH/H2 regions when ^15^N-labeled IC_1-88_ was titrated with unlabeled IC_100-260_ (Fig 5C).

### Interactions of p150_CC1B_ and NudE_CC_ with CT IC

The interaction between CT IC_1-88_ (a smaller IC construct IC_1-88_ that includes only the p150 and NudE binding domains) and p150_CC1B_ has been previously reported (22) and is shown here only for comparative purposes (Fig 6C, top left). Here, we explore the interaction between IC_1-88_ and NudE_CC_ under similar conditions. By SV-AUC, IC_1-88_ bound to p150_CC1B_ shows a sedimentation coefficient of 3.7, whereas for IC_1-88_ bound to NudE_CC_ the coefficient is only 2.7 (Fig 6A). This is surprising considering that the two complexes have similar overall masses and binding stoichiometries, and suggests that the complex with NudE_CC_ is less compact than the complex with p150_CC1B_. ITC indicates that, like IC_1-88_ and p150_CC1B_ (Fig 6C, top left), IC_1-88_ and NudE_CC_ bind with a 1:1 molar ratio (Fig 6C, bottom left), which corresponds to one NudE_CC_ dimer binding two IC_1-88_ monomeric chains. IC_1-88_ binds to NudE_CC_ with a dissociation constant (*K*_d_) of 0.3 μM, which is weaker than the nM affinity estimated for IC_1-88_ binding to p150_CC1B_ (22). Previously published NMR titrations of unlabeled p150_CC1B_ with ^15^N-labeled IC constructs that contain (IC_1-88_) or do not contain (IC_37-88_) the SAH region demonstrate that both the SAH and H2 regions bind to p150_CC1B_ (22) (Fig 6B). IC_1-88_ binding to p150_CC1B_ exhibits an unusual two-step ITC thermogram (Fig 7C, top left) whereas a one-step thermogram is seen when NudE_CC_ is titrated with IC_1-88_ (Fig 7C, bottom left), suggesting a difference in the mode of binding. We propose that the two step thermogram is due to binding to both the SAH and H2 regions of IC_1-88_ with p150_CC1B_, while in contrast the single step thermogram is due to NudE binding only to the SAH region.

**Figure 6.**
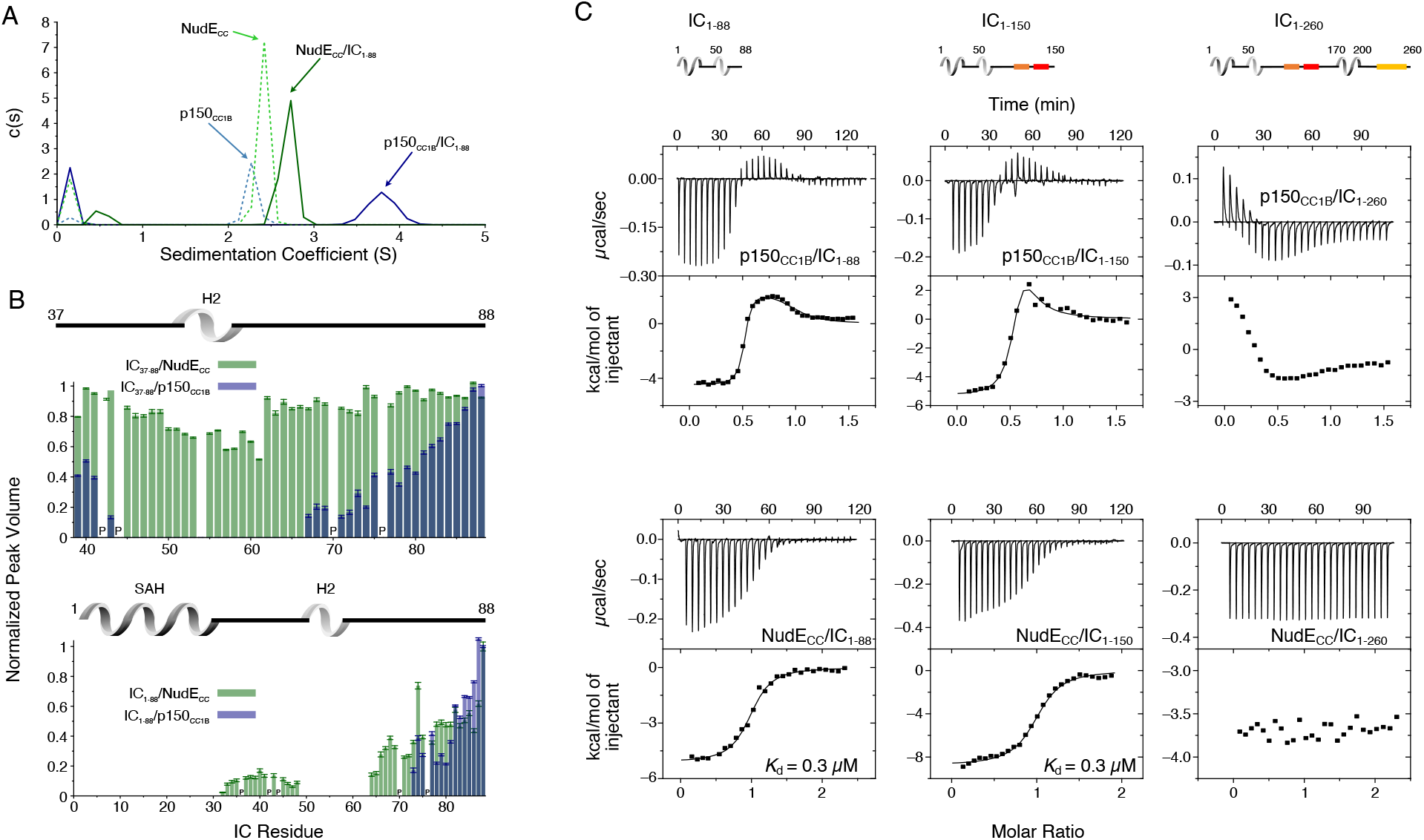
Binding interactions of CT IC to p150_CC1B_ and NudE_CC_. (A) SV-AUC profiles for samples containing p150_CC1B_ (blue dashed line), NudE_CC_ (green dashed line), IC_1-88_/p150_CC1B_ complex (blue solid line) and IC_1-88_/NudE_CC_ complex (green solid line) show that IC_1-88_ complexes have a larger sedimentation coefficient with p150_CC1B_ than with NudE_CC_. No data were collected for free IC_1-88_ because it has no absorbance at 280 nm. (B) Normalized peak volumes in the ^1^H-^15^N HSQC spectra for ^15^N-labeled IC_37–88_ (top) or ^15^N-labeled IC_1–88_ (bottom) when titrated with unlabeled p150_CC1B_ (blue) and NudE_CC_ (green). “P” indicates proline residues. No peak disappearance for IC_37-88_ was observed when NudE_CC_ was added. (C) ITC thermograms for p150_CC1B_ titrated with IC_1-88_ (top left), IC_1-150_ (top middle), and IC_1-260_ (top right), and for NudE_CC_ titrated with IC_1-88_ (bottom left), IC_1-150_ (bottom middle), and IC_1-260_ (bottom right), collected at 25°C (pH 7.5). For IC_1-260_, reduced and endothermic binding is observed with p150_CC1B_ whereas no binding is observed with NudE_CC_. All results indicate that the H2 region of CT IC binds weakly to p150_CC1B_ but does not bind NudE_CC_. **Figure 6-source data 5: Source files for SV-AUC and NMR binding studies of IC fragments with p150_CC1B_ and NudE_CC_.** This Excel workbook contains the data plotted for the SV-AUC experiments as well as the NMR peak volume ratios for binding studies between ^15^N-labeled IC_1-88_ and ^15^N-labeled IC_37-88_ with p150_CC1B_ and NudE_CC_. Additional information regarding data collection can be found in the corresponding methods section. Data were plotted using gnuplot.

**Figure 7.**
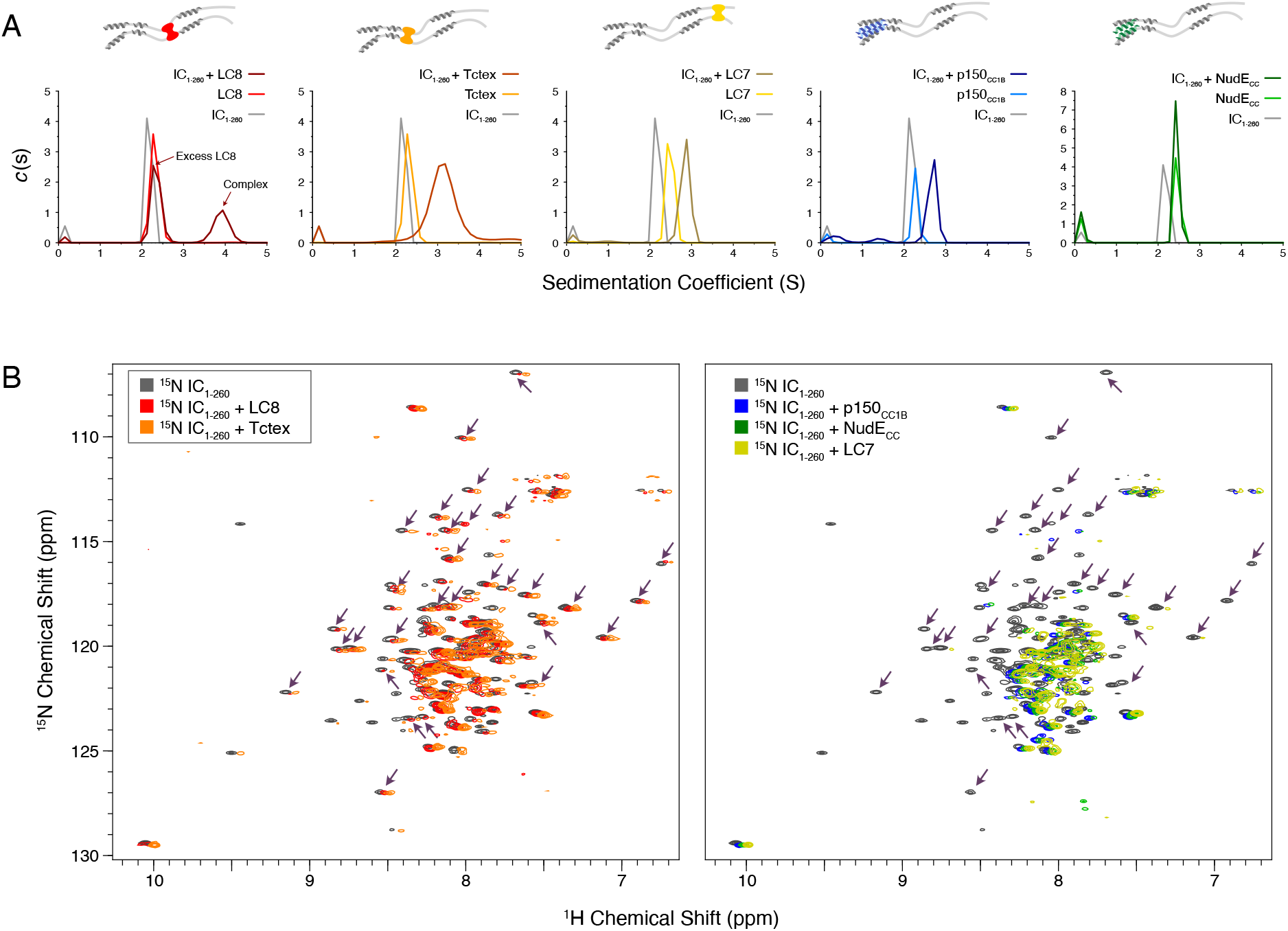
Binding characterization of binary complexes of IC_1-260_. (A) SV-AUC of IC_1-260_/LC8, IC_1-260_/Tctex, IC_1-260_/LC7, IC_1-260_/p150_CC1B_, and IC_1-260_/NudE_CC_. Data for the binary complexes is overlayed with data for each protein individually to better see shifts in the sedimentation coefficient of the binary complexes. (B) ^1^H-^15^N TROSY overlays of free ^15^N-labeled IC_1-260_ (black) and ^15^N-labeled IC_1-260_ bound to unlabeled binding partners in a 1:1.5 molar ratio. The spectra were offset by 0.03 ppm in the ^1^H dimension to help illustrate changes in peak intensities. Changes in peak appearances/shifts/disappearances seem to be similar for LC8 (red) and Tctex (orange) versus changes seen for p150_CC1B_ (blue), NudE_CC_ (green), and LC7 (yellow). Arrows indicate peaks that remain when LC8 and Tctex are added to IC_1-260_, but disappear when p150_CC1B_, NudE_CC_, and LC7 are added. **Figure 7-source data 6: Source files for SV-AUC binding studies of IC_1-260_.** This Excel workbook contains the data plotted for the SV-AUC experiments shown in Figure 7. Additional information regarding data collection can be found in the corresponding methods section. Data were plotted using gnuplot.

To identify if NudE_CC_ does indeed only bind to the SAH region or to both the SAH and the H2 regions, NMR titrations with unlabeled NudE_CC_ and ^15^N-labeled IC_1-88_ or IC_37-88_ constructs were carried out. In contrast to the NMR titration with p150_CC1B_, peaks for residues corresponding to the H2 region did not disappear upon titration of ^15^N IC_37-88_ with NudE_CC_ (Fig 6B, top), confirming our proposal that the IC H2 region does not directly bind to NudE_CC_ whereas this region does directly bind to p150_CC1B_. For both p150_CC1B_ and NudE_CC_, titration into ^15^N IC_1-88_ resulted in IC_1-88_ peaks disappearing for both the SAH and H2 regions (Fig 6B, bottom). The disappearance of IC_1-88_ peaks for both the SAH and H2 regions with NudE_CC_ can be explained by an interaction between the SAH and H2 regions (17) that relays the change in correlation time of the SAH region upon binding to NudE_CC_ to the H2 region. The difference in how IC binds p150_CC1B_ and NudE_CC_ observed in the ITC experiment is conserved when a somewhat longer IC construct is used (IC_1-150_, Fig 6C, center), whereas binding is severely diminished when using a construct containing the entire N-IC region (IC_1-260_, Fig 6C, right). Further data and discussion of these observations are presented below.

### Interactions using CT IC_1-260_

SV-AUC was initially used to characterize complex formation between IC_1-260_ and each of the other binding partners. The largest sedimentation coefficient was observed for the IC_1-260_/LC8 complex; for all other complexes, a less dramatic peak shift was seen, a possible indicator of weaker binding, a dynamic equilibrium between free and bound states, and/or a shift to a more elongated conformation (Fig 7A). For the IC_1-260_/NudE_CC_ complex, no significant change was observed in the SV-AUC data relative to unbound NudE_CC_. However, the absence of a peak corresponding to free IC for the sample containing IC_1-260_ and NudE_CC_ indicates that some degree of binding takes place. This result along with our ITC (Fig 6C, bottom right) and NMR (Fig 7B, right) data for this complex indicates that binding of IC_1-260_ to NudE_CC_ is very weak and is thus only observed at high protein concentrations.

Although we were unable to assign the NMR spectrum of IC_1-260_ at 40°C because of poor sensitivity and unfavorable relaxation times, overlays of ^1^H-^15^N TROSY spectra for free ^15^N-labeled IC_1-260_ and binary complexes of ^15^N-labeled IC_1-260_ with unlabeled binding partners (Fig 7B) show distinct patterns of peak disappearances. Upon addition of Tctex or LC8 (Fig 7B, left), only a handful of IC_1-260_ peaks disappear and the patterns of disappearances are similar, as expected based on the proximity of the Tctex and LC8 binding sites. When either LC7, p150_CC1B_, or NudE_CC_ is added, considerably more peaks disappear in the spectra for ^15^N-labeled IC_1-260_ (Fig 7B, right) and, surprisingly, similar patterns of disappearances occur even though the LC7 binding site is at the C-terminus of IC_1-260_ whereas the p150_CC1B_ and NudE_CC_ binding sites are at the N-terminus. The similar patterns of peak disappearances suggests that regions of the N and C-termini of IC_1-260_ interact in such a way that when one end of IC_1-260_ is bound it affects the peak intensities of the other end and vice versa.

### Multivalency relieves IC autoinhibition

Following characterization of individual binding events, we sought to reconstitute full N-IC subcomplexes. Each individual subunit was first expressed and purified individually prior to dynein subcomplex formation (IC_1-260_/Tctex/LC8/LC7), achieved by mixing IC_1-260_ with the dynein light chains in a 1:1.5 (IC to LC) molar ratio. To this dynein subcomplex, p150_CC1B_ or NudE_CC_ was added to create two larger subcomplexes. Each subcomplex was re-purified by size exclusion chromatography (SEC) to remove any excess of the binding partners (which elute at ∼215 mL) and to assess their overall stability (Fig 8A). The shape and symmetry of the eluting SEC peak for the dynein subcomplex (∼140 mL) indicates a weaker binding affinity than when either p150_CC1B_ (∼120 mL) or NudE_CC_ (∼130 mL) was added. In all cases however, each expected subunit was detected by sodium dodecyl sulphate–polyacrylamide gel electrophoresis (SDS-PAGE) of collected fractions (Fig 8B), thus validating successful assembly.

**Figure 8:**
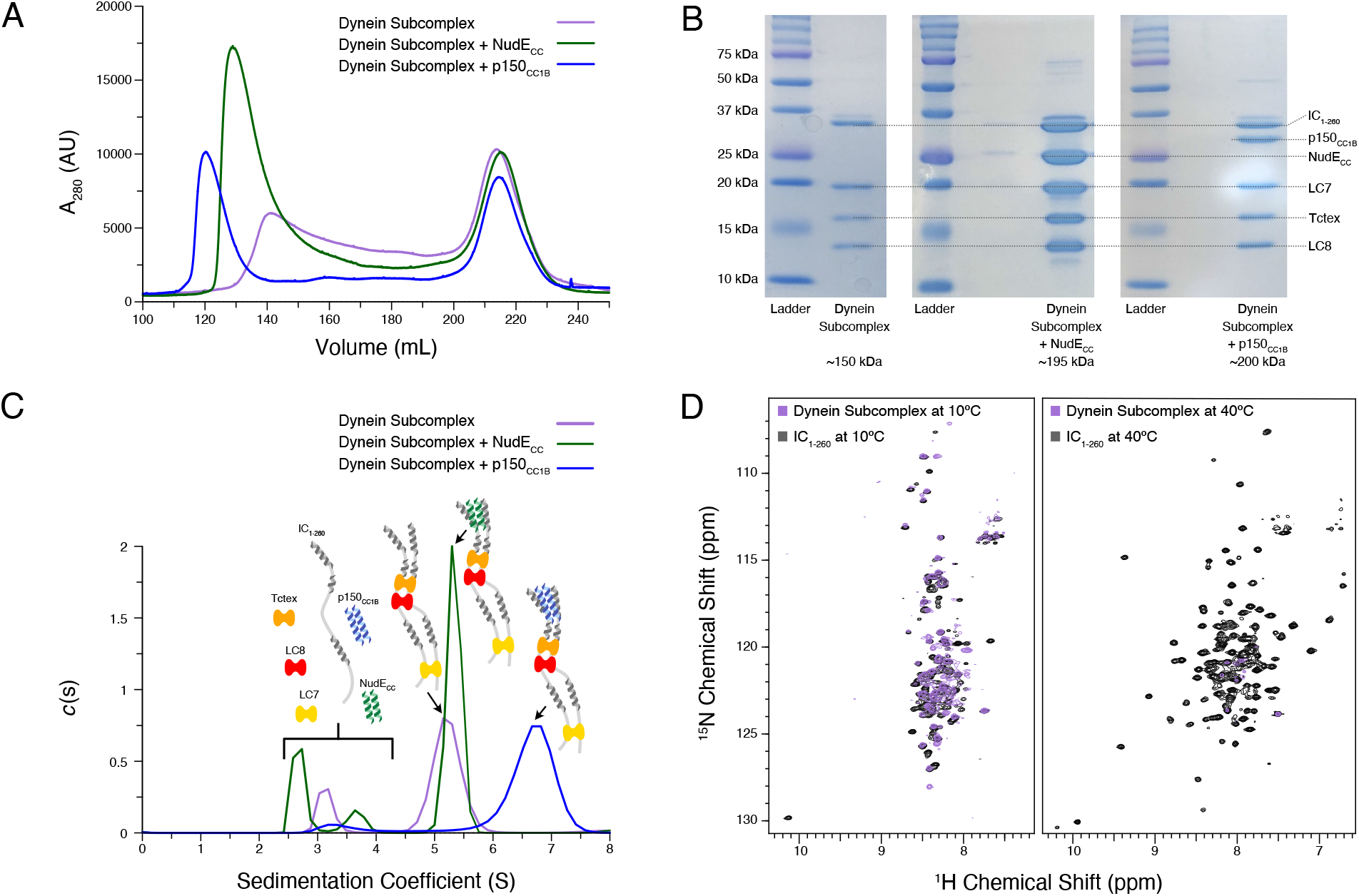
Reconstitution and characterization of dynein subcomplexes. (A) SEC traces of the dynein subcomplex (IC/light chains) (purple) and the dynein subcomplex with the addition of either p150_CC1B_ (blue) or NudE_CC_ (green). (B) SDS-PAGE gels of fractions collected from SEC for all complexes showing all expected proteins. (C) SV-AUC profiles of the dynein subcomplex (purple) bound to p150_CC1B_ (blue) or NudE_CC_ (green). (D) ^1^H-^15^N TROSY overlays of free IC_1-260_ (black) and the dynein subcomplex (purple). At 10°C many peaks are still of high intensity in the 153 kDa complex, indicating that some regions remain disordered. The very few peaks at 40°C of the bound is most likely due to the size and tumbling of the subcomplex and consistent with the fact that the majority of the peaks at this temperature are from ordered regions. **Figure 8-source data 7: Source files for SEC and SV-AUC of reconstituted IC_1-260_ subcomplexes.** This Excel workbook contains the data plotted for the SEC and SV-AUC experiments shown in Figure 8 with individual sheets corresponding to panels within the figure. Additional information regarding data collection can be found in the corresponding methods section. Data were plotted using gnuplot. **Figure 8-source data 8: Original SDS-PAGE gel images.** This zipped folder contains the original files of the full raw unedited gel images for the SEC purification of each IC_1-260_ subcomplex. There is also a combined image with the uncropped gels with the relevant bands clearly labeled.

Using SV-AUC, we show that when all dynein light and intermediate chains are present, the fully bound dynein subcomplex sediments as a single peak with a sedimentation coefficient of approximately 5 S (Fig 8C). Surprisingly, addition of NudE_CC_, which adds approximately 45 kDa of mass, only slightly increases the sedimentation coefficient. In contrast, addition of p150_CC1B_ increases the sedimentation coefficient to approximately 7 S. The difference in the SV-AUC data between the addition of p150_CC1B_ and NudE_CC_ is surprising based on the similar expected masses of the complexes (201 and 198 kDa, respectively). This difference may be explained by the overall shape of the two bound complexes, details of which need to be further examined. Also, important to note is that the shift in sedimentation coefficient for p150_CC1B_ when added to the IC/light chains complex is significantly more pronounced than that with IC_1-260_ alone, suggesting that binding to IC_1-260_ is significantly enhanced in the presence of the light chains.

NMR spectroscopy indicates that most of the peaks in the disordered regions observed at 10°C for free IC_1-260_ (Fig 8D) remain in the spectrum for the dynein subcomplex (IC_1-260_/Tctex/LC8/LC7), indicating that there is still significant disorder in the fully bound complex (153 kDa). In contrast, comparison of the spectra of free IC_1-260_ and the dynein subcomplex at 40°C shows a drastic disappearance of peaks, indicating that the residues corresponding to these peaks have much longer correlation times in the bound state either due to their involvement in binding or due to the overall increase in weight of the entire subcomplex. Peaks from linker regions not involved in binding light chains that are observed at 10°C but are not observed at 40°C likely disappear due to rapid exchange with the solvent at the higher temperature.

### Autoinhibition is retained in full-length CT IC

To determine if the autoinhibition seen in IC_1-260_ is also present in the full-length construct, IC_FL_ (residues 1-642) was produced using a baculovirus expression system. The estimated mass of 75.4 kDa for IC_FL_ from SEC-MALS matches closely to the expected monomeric mass of 79 kDa (Fig 9A-B). Further, SV-AUC shows a single, homogenous peak with a sedimentation coefficient of 4.0 S, as would be expected for monomeric IC_FL_ (Fig 9B top). This is the first study that shows full-length IC is a monomer in solution and requires the light chains for its dimerization. Adding either p150_CC1B_ or NudE_CC_ to IC_FL_ results in a negligible shift of the IC_FL_ peak and, in both cases, a peak corresponding to unbound p150_CC1B_ or NudE_CC_ was observed (Fig 9B). This lack of binding between IC_FL_ and p150_CC1B_/NudE_CC_ shows that autoinhibition occurs in IC_FL_ in a similar manner to what we have already observed in the IC_1-260_ construct. SV-AUC experiments on the dynein subcomplex interactions confirm that the autoinhibited state is released by the addition of the light chains. Full length IC bound to the three light chains (IC_FL_/Tctex/LC8/LC7) has a sedimentation coefficient of 7.0 S but shifts to values of 7.5 and 8 S upon addition of NudE_CC_ and p150_CC1B_, respectively (Fig 9B middle). These results mimic those seen for the IC_1-260_ construct and indicate that the addition of the light chains allows p150_CC1B_ or NudE_CC_ to bind by relieving IC autoinhibition.

**Figure 9.**
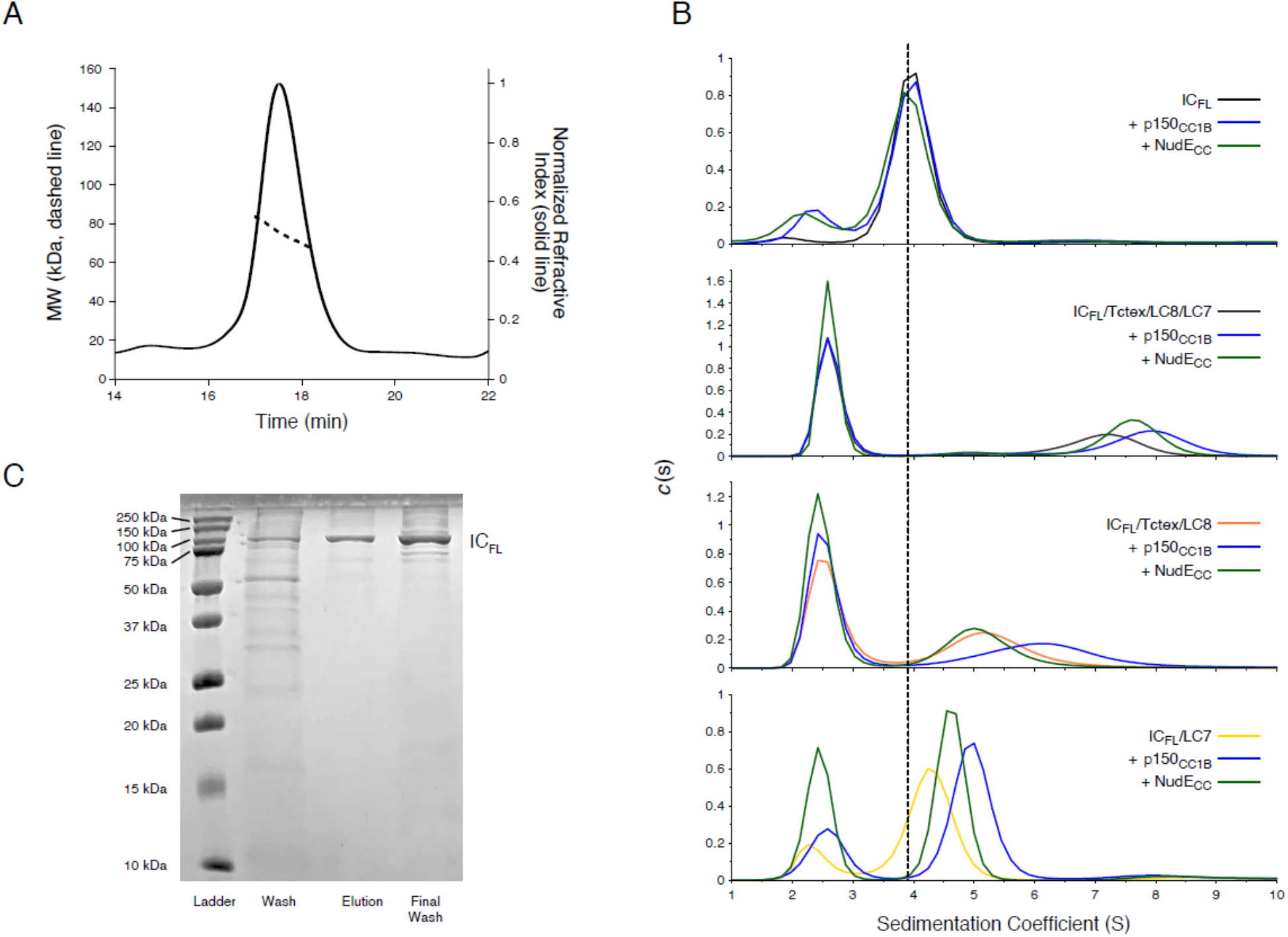
Binding characterization of CT IC_FL_ subcomplexes. (A) The estimated mass of IC_FL_ from MALS is 75.4 kDa, which indicates that IC_FL_ is a monomer in the absence of binding partners. (B) SV-AUC profiles of IC_FL_ (black), IC_FL_ mixed with p150_CC1B_ (blue) or NudE_CC_ (green), and the subcomplexes: IC_FL_/Tctex/LC8/LC7 (gray), IC_FL_/Tctex/LC8/LC7/p150_CC1B_ (blue), IC_FL_/Tctex/LC8/LC7/NudE_CC_ (green), IC_FL_/Tctex/LC8 (orange), IC_FL_/Tctex/LC8/p150_CC1B_ (blue), IC_FL_/Tctex/LC8/NudE_CC_ (green), IC_FL_/LC7 (yellow), IC_FL_/LC7/p150_CC1B_ (blue), and IC_FL_/LC7/NudE_CC_ (green). The black, dashed line is centered on unbound IC_FL_ to help guide the eye. (C) SDS-PAGE gel of IMAC fractions (left to right: wash, elution, and final wash) with a band for IC_FL_ migrating in accordance with the expected mass of ∼79 kDa. **Figure 9-source data 9: Source files for SEC-MALS and SV-AUC of IC_FL_.** This Excel workbook contains the data plotted for the SEC-MALS experiment of IC_FL_ and the SV-AUC binding experiments shown in Figure 9 with individual sheets corresponding to panels within the figure. Additional information regarding data collection can be found in the corresponding methods section. Data were plotted using gnuplot. **Figure 9-source data 10: Original SDS-PAGE gel image.** This zipped folder contains the original file of the full raw unedited gel image for the IMAC fractions of IC_FL_. There is also an image with the uncropped gel with the relevant bands clearly labeled. Notice, two batches of IC_FL_ were purified in tandem.

To further explore the role of each of the light chains in releasing IC autoinhibition, SV-AUC experiments with IC_FL_/Tctex/LC8 and IC_FL_/LC7 were conducted. The IC_FL_/Tctex/LC8 and IC_FL_/LC7 complexes show peaks with sedimentation coefficients of approximately 5.2 and 4.2 S, respectively (Fig 9B). Interestingly, the IC_FL_/Tctex/LC8 complex exhibits no shift upon the addition of NudE_CC_ but does shift to 6.2 S upon the addition of p150_CC1B_ (Fig 9B). The IC_FL_/LC7 complex, on the other hand, exhibits a shift upon addition of either NudE_CC_ or p150_CC1B_ to 4.5 or 5 S, respectively (Fig 9B). These data suggest that the addition of LC7 is sufficient to allow binding of p150_CC1B_ and NudE_CC_ to IC_FL_, whereas the binding of Tctex and LC8 to IC_FL_ only promotes p150_CC1B_ binding to IC_FL_.

## Discussion

The N-terminus of IC from a variety of species contains a stretch of about 300 amino acids that are primarily disordered, except for a few short α-helices. Within the first 40 residues is a fully ordered helix (the SAH region), followed by a short disordered linker and another region (H2) that forms a fully ordered helix in some species but is only a nascent helix in others (3,20,22,37). Prior work suggests that a more disordered H2 is correlated with tighter IC/p150^Glued^ binding and has shown that, for CT IC, the H2 region binds directly to p150^Glued^ (although mostly in a nonspecific manner) (22). In this work, we use a construct of IC that encompasses almost its entire 300-amino acid N-terminus to probe the interactions of CT IC with both p150_CC1B_ and NudE_CC_. This construct allows us to study the assembly of IC into a multivalent subcomplex with three dimeric dynein light chains and with its binding partners, p150^Glued^ and NudE. Further, we demonstrate, using full-length IC, that the mechanisms at place in IC_1-260_ remain in the context of the entire IC protein. A model that illustrates the importance of autoinhibition in dynein regulation and how the assembly of the multivalent IC subcomplex relieves this autoinhibition is presented (Fig 10).

**Figure 10:**
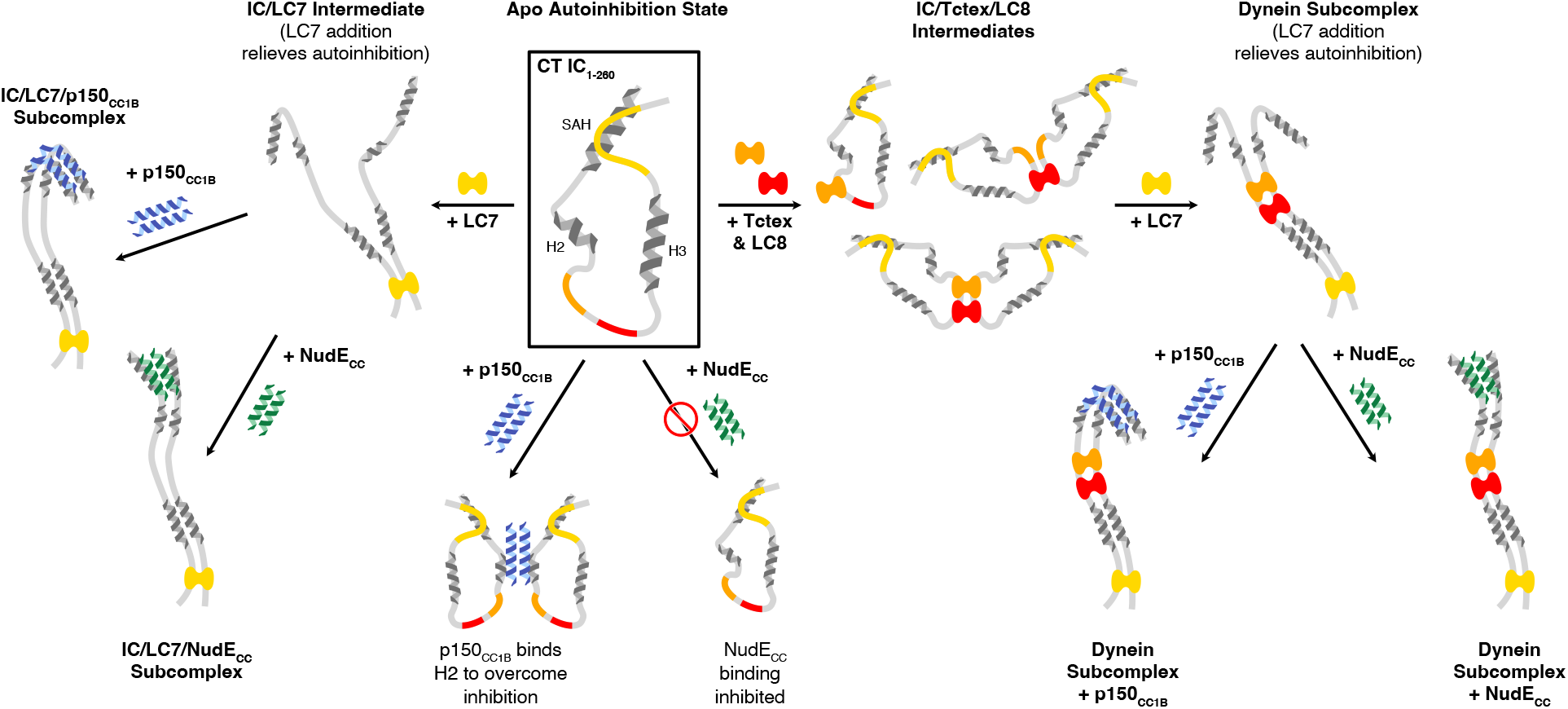
A model of CT IC_1-260_ binding interactions and subcomplex assemblies. Apo CT IC_1-260_ is compact and in autoinhibited state (Boxed in black), with the SAH, H2, H3 regions depicted as helices and with colors indicating the LC8 (red), Tctex (orange), and LC7 (yellow) binding sites. When LC7 is added (left arrow) LC7 outcompetes autoinhibition to bind IC_1-260_, exposing the SAH domain for p150_CC1B_ and NudE binding. When p150_CC1B_ or NudE_CC_ are added (down arrows) to apo IC_1-260_, autoinhibition prevents NudE_CC_ from binding and reduces the binding affinity of p150_CC1B_. However, as p150_CC1B_ is able to bind to the H2 region of IC_1-260_, binding is not completely prevented. Addition of Tctex and/or LC8 (right arrow) leads to a number of possible binary and ternary intermediates. LC8’s role of driving IC dimerization is depicted and, in all intermediates, we predict that IC autoinhibition remains based on very limited changes in NMR spectra. Continuing to the right, the addition of LC7 leads to formation of the dynein subcomplex and the release of SAH autoinhibition. The free SAH is now able to resume transient interactions with H2 prior to binding with either p150_CC1B_ or NudE_CC_. Finally, addition of p150_CC1B_ or NudE_CC_ leads to the formation of the p150 and NudE subcomplexes (bottom right). Our SV-AUC data suggest that the NudE subcomplex adopts a more elongated conformation.

### CT IC_1-260_ is a partially disordered compact monomer

The primary CT IC construct (IC_1-260_) used in this work is, to date, the longest IC construct from any species made recombinantly and extensively studied by NMR, ITC, and SV-AUC (2,3,12,14,20,21,30,31,37). IC_1-260_ far exceeds previously studied constructs from CT (res. 1-35, 37-88, and 1-88) as well as constructs from Drosophila (res. 1-60, 30-143, 84-143, 1-143, 92-260, and 114-260) and rat (res. 1-44, 1-96, and 1-112). Similar to IC from Drosophila, apo CT IC_1-260_ is monomeric (Fig 2A, 2D) and contains significant disorder (Fig 4). However, unlike ICs from other species, CT IC_1-260_ has a sequence-predicted third helix, H3, corresponding to residues 170-200 (Fig 3A, S2) that is highly consistent with fractional helicity values of IC_1-260_, IC_160-240_, and IC_100-260_ determined by CD spectroscopy (Fig 3B-E). IC_1-260_ is the most thermostable construct, resisting unfolding at temperatures of up to 50°C (Fig 3B), a feature that we attribute to long-range tertiary contacts between the N- and C-termini (Fig 5, S3). SV-AUC and SEC-MALS experiments confirm the presence of a binding interaction between IC_1-88_ and IC_100-260_ (Fig S3). Further, NMR titrations with N- and C-terminal IC constructs identify a self-association interaction between residues in the SAH and the LC7 binding site of IC_1-260_ (Fig 5). These tertiary contacts are also seen in comparisons of NMR spectra of ^15^N-labeled IC_1-260_ upon addition of p150_CC1B_, NudE_CC_, or LC7, which reveal a similar pattern of peak disappearances and chemical shifts that are not the same as those observed upon addition of Tctex or LC8 (Fig 7B). Finally, long distance tertiary interactions, and their impact on IC binding interactions, are suggested by the dramatic reduction in binding of p150_CC1B_ or NudE_CC_ with IC_1-260_ compared to IC_1-88_ and IC_1-150_ (Fig 6C).

### Different modes of binding of IC to p150^Glued^ and NudE

The coiled-coil domains of p150^Glued^ and NudE (p150_CC1B_ residues 478-680, and NudE_CC_ residues 1-190) show similar structure propensity and stability by SV-AUC and CD and have stable dimeric structures with dissociation constants of 0.03 and 0.20 µM, respectively (Fig 2). However, despite their many similarities, p150_CC1B_ and NudE_CC_ have different modes of binding to CT IC. First, IC interactions with p150_CC1B_ are multistep, which we attribute to p150_CC1B_ binding to both the SAH and H2 regions of IC (22). In contrast, NudE_CC_ binds IC in a single step (Fig 6C) and only to the SAH region and not to the H2 region (Fig 6B). Second, the p150_CC1B_/IC_1-88_ complex has a larger sedimentation coefficient when compared to the NudE_CC_/IC_1-88_ complex (3.8 S vs 2.7 S, Fig 6A) even though the complexes have similar expected masses and ITC shows that they have the same binding stoichiometry. This suggests that the p150_CC1B_/IC_1-88_ complex has a more compact structure. Third, although binding is much weaker with IC_1-260_ (Fig 6C) compared to the smaller IC constructs for both p150_CC1B_ and NudE_CC_, binding to NudE_CC_ is significantly weaker and is only detected at the higher concentrations used in NMR experiments (Fig 7B). Finally, and most importantly, the light chain/IC assembled complex shows a different sedimentation coefficient when bound to p150_CC1B_ or NudE_CC_. The subcomplexes both have masses of almost 200 kDa, but the NudE_CC_ subcomplex has a sedimentation coefficient of ∼5 S compared to ∼7 S for the p150_CC1B_ subcomplex (Fig 8C). The larger sedimentation coefficient for the p150_CC1B_ subcomplex is unexplained by the small mass difference, and thus indicates a more compact and stable conformation for the subcomplex when p150_CC1B_ is bound. While adding p150_CC1B_ makes the light chain/IC assembled complex heavier and increases the sedimentation coefficient, adding NudE_CC_ does not result in a significant increase, suggesting both weaker binding and increased friction compared to the light chain/IC/ p150_CC1B_ assembled complex. A difference in overall complex mass due to a binding stoichiometry greater than 1:1 for p150_CC1B_ and IC could explain the complex’s larger sedimentation coefficient, however this is very unlikely. As shown in previous studies across multiple species, p150_CC1B_ consistently binds IC in a 1:1 stoichiometry (2,17,20–22). Furthermore, visualization of the light chain/IC/p150_CC1B_ complex and the light chain/IC/NudE_CC_ complex by SDS-PAGE (Fig 8B) shows similar intensity ratios between IC_1-260_/NudE_CC_ and IC_1-260_/p150_CC1B_, indicating that they exist in the same stoichiometry. Finally, MALS data of the IC_1-88_/p150_CC1B_ complex gives a mass of 66 kDa, consistent with the mass expected for a p150_CC1B_ dimer and two monomeric IC_1-88_ chains (data not shown). SV-AUC experiments with IC_FL_ further confirm that p150_CC1B_ and NudE_CC_ have different binding modes as, once again, complexes with assembled IC_FL_/light chain subcomplexes show a larger sedimentation coefficient when bound to p150_CC1B_ (∼8 S) than when bound to NudE_CC_ (∼7.5 S) (Fig 9).

### Autoinhibition in IC selects for binding of p150^Glued^ over NudE

Interactions of IC_1-260_ with its binding partners observed by ITC and AUC show weak endothermic binding to p150_CC1B_ but no binding to NudE_CC_. Our data suggest a process, which we refer to as autoinhibition, in which IC_1-260_ adopts a compact structure that covers the SAH region necessary for binding to NudE_CC_ and for strong binding to p150_CC1B_, but still leaves the H2 region partly accessible so that weak binding to p150_CC1B_ can still occur. The SAH region is made inaccessible by tertiary interactions within IC, as seen in an NMR titration of ^15^N-labeled IC_1-88_ with IC_100-260_ (Fig 5B-C). We assign these tertiary interactions to those within the LC7 binding site at the C-terminus via NMR titrations of ^15^N-labeled IC_216-260_ with IC_1-88_ (Fig 5D-E) but note that the binding affinity does appear stronger in the longer construct (IC_100-260_) (Fig S3, 5B-C) indicating that the full context of the disordered chain encompassing the binding sites for Tctex, LC8, and LC7 is needed for strong, intramolecular interactions. These interactions likely have an autoinhibitory effect, which would explain the reduced binding affinity for p150_CC1B_ and NudE_CC_ for IC_1-260_ in comparison to the shorter constructs. Since this effect is more pronounced for IC_1-260_ binding with NudE_CC_, we propose autoinhibition as a mechanism for partial binding of IC_1-260_ with p150_CC1B_ and thus selection of p150_CC1B_ over NudE_CC_. Autoinhibition as a selection mechanism is underscored by our results with full-length IC (Fig 9). Although binding to both p150_CC1B_ and NudE_CC_ are inhibited with IC_FL_ alone, the addition of Tctex and LC8 rescue binding to p150_CC1B_, likely by helping make the nearby H2 region more available. In contrast, binding to both p150_CC1B_ and NudE_CC_ is rescued via LC7 binding to IC_FL_ (Fig 9).

### Regulation of binding to p150^Glued^ across species and the role of dynein light chains

Based on current data, it is highly likely that autoinhibition in IC is a conserved process for modulating binding to p150^Glued^ and NudE. In Drosophila IC, paramagnetic relaxation enhancement NMR experiments have shown that binding of NudE to the SAH region shifts IC equilibrium toward more open states (17). Such a preference for more open states when NudE is bound is in line with our data for CT showing that NudE_CC_ requires an open SAH region to bind, whereas p150_CC1B_ can more easily overcome the autoinhibited (closed) conformation of IC. Similarly, with IC from mammalian species, autoinhibition by interaction between the SAH and H2 regions appears to be modulated by phosphorylation (20, 54). Our data collected on the full N-terminal domain of IC and full-length IC protein from CT demonstrate subcomplex assembly via light chain binding as the primary modulating mechanism for p150^Glued^ and NudE binding. This is an effect that can only be detected with larger constructs of IC, such as CT IC_1-260_, that contains long disordered linkers separating short helices and binding sites. As summarized by the model shown in Figure 10, we propose that these properties combine to create a multifaceted system of regulation for IC. The disordered linkers allow for the needed flexibility to bring together the SAH region and the C-terminus (200+ amino acids away) and to extend it into an open conformation when LC7 is bound (either on its own, or with the other two light chains), which suggests a unique role for LC7 in regulating IC interactions.

In conclusion, this work shows, for the first time, the recombinant expression of multiple dynein and non-dynein subunits and their reconstitution into three assembled subcomplexes. The dynein subcomplex (IC/Tctex/LC8/LC7) as well as the dynein subcomplex with p150_CC1B_ or with NudE_CC_ were each successfully re-purified by SEC (Fig 8A) and verified by SDS-PAGE to show each expected subunit present in the complex (Fig 8B). Both the shape and the symmetry of the eluting SEC peak for the dynein subcomplex indicate that the complex is less stable than when either p150_CC1B_ or NudE_CC_ is added. Taken together, these data demonstrate that not only is the stability of the dynein subcomplex enhanced by addition of p150_CC1B_ and NudE_CC_, but also that the ability of p150_CC1B_ and NudE_CC_ to bind to IC is no longer inhibited when the light chains are present. The same conclusion is supported by data with full-length IC, as IC_FL_ alone is unable to bind to either p150_CC1B_ or NudE_CC_ until light chains are bound (Fig 9). Interestingly, the binding of just LC7 is enough to rescue IC_FL_ binding to p150_CC1B_ and NudE_CC_, while binding of Tctex and LC8 only relieves autoinhibition enough for p150_CC1B_ to bind (Fig 9B). These results speak to the complexity required in IC regulation and emphasize the roles of autoinhibition and multivalency in this process.

As illustrated in the model in Figure 10, the C-terminal end of IC_1-260_, which contains the LC7 binding site, interacts with the SAH region and results in a closed conformation. The closed conformation of IC is autoinhibitory, preventing binding of the SAH region by p150_CC1B_ or NudE_CC_. However, because p150_CC1B_ also binds to the H2 region, p150_CC1B_ binding to IC_1-260_ is not completely abolished. Our data suggest that it is the segment containing residues 220-250 (which forms an α-helix upon binding LC7) that makes contacts with the SAH, leaving downstream residues able to initiate binding to LC7 (Fig 5, 9). From the limited changes in NMR spectra upon Tctex and LC8 binding, we conclude that the overall closed conformation of IC is likely unrelieved by binding of LC8 and/or Tctex since they bind in the center of a long linker, whereas LC7 binding is required to release the SAH region, thereby allowing assembled IC to fully bind to p150_CC1B_ and NudE_CC_.

Dynein autoinhibition has been a relevant topic for years as dynein is only weakly processive when isolated in vitro (55), and requires the binding of dynactin and/or cargo adaptor proteins for activation (56–59). In a recent study, cryo-EM structures revealed that the autoinhibited form of human cytoplasmic dynein (phi-particle) is stabilized by motor domain self-dimerization and contact between the tails of the heavy chains, whereas the less inhibited form (open-dynein) formed after binding dynactin has the proper motor domain orientation for highly processive motion along the microtubules (7). Due to the binding relationship between the N-terminus of IC and the CC1B region of the p150^Glued^ subunit of dynactin, there is the possibility that the IC/p150^Glued^ interaction provides the first step in dynein activation. If so, it would stand to reason that IC/p150^Glued^ autoinhibition regulation we identify here in CT is a critical step in dynein function. In future work it will be important to demonstrate that autoinhibition and its subsequent reversal by dynein light chain binding is a conserved process of IC across species and whether this plays a role in the autoinhibition of the overall dynein protein complex.

## Methods

### Cloning, Protein Expression and Purification

All studies were carried out using constructs from *Chaetomium thermophilum* (thermophilic fungus, G0SCF1-1). IC_1-260_ (res. 1-260), p150_CC1B_ (res. 478-680), and NudE_CC_ (res. 1-190) constructs were prepared by PCR and cloned into a pET-24d vector with an N-terminal 6×His tag using the Gibson Assembly protocol (60, 61). In addition, a fragment of the IC_1-260_ construct with N-terminal residues removed (IC_100-260_) was generated by the same method. IC_160-240_ was ordered from GenScript (Piscataway, NJ) and IC_216-260_ was ordered from Azenta Life Sciences (Chelmsford, MA). Full length CT Tctex, LC8, and LC7 were amplified out of a CT cDNA library and cloned into a pET-15b vector with an uncleavable C-terminal 6×His tag. These light chain constructs contain a single, non-native GS linker prior to the tag sequence. For IC, p150, and NudE constructs, an N-terminal tobacco etch virus (TEV) protease cleavage site was included to allow the removal of the 6×His tag, leaving a non-native GAH sequence post cleavage. DNA sequences were verified by Sanger sequencing. IC_1-88_ and IC_37-88_ constructs were prepared previously (22).

Recombinant plasmids were transformed into Rosetta (DE3) *E. coli* cells (Merck KGaA, Darmstadt, Germany) for protein expression. Bacterial cultures for expression of unlabeled proteins were grown in ZYM-5052 autoinduction media at 37°C for 24 hrs (62), whereas cultures for expression of isotopically-labeled (^15^N or ^15^N/^13^C) proteins were grown in MJ9 minimal media (63) at 37°C to an OD_600_ of 0.8 before being induced with 0.4 mM isopropyl-β-D-1-thiogalactopyranoside (IPTG), and continuing expression overnight at 26°C. Proteins were purified from the cell cultures by immobilized metal affinity chromatography using previously published methods (22). Complete cleavage of the tag by tobacco etch virus protease was verified by SDS-PAGE analysis.

Proteins were further purified using a Superdex 75 (Cytiva Life Sciences, Pittsburgh, PA) SEC column and then, for IC_1-260_ samples, followed by anion exchange using Macro-Prep High Q Support resin (Bio-Rad, Hercules, California) with elution in 0.1 to 0.2 M sodium chloride. Protein concentrations were determined from absorbance at 205 and 280 nm (64). Molar extinction coefficients for the constructs used are as follows (ε_205_ & ε_280_): IC_1-260_ = 853,230 & 11,460 M^-1^ cm^-1^, IC_1-150_ = 488,290 & 8,480 M^-1^ cm^-1^, IC_100-260_ = 665,620 & 14,440 M^-1^ cm^-1^, IC_160-240_ = 387,110 & 2,980 M^-1^ cm^-1^, p150_CC1B_ = 685,820 & 9,970 M^-1^ cm^-1^, NudE_CC_ = 665,800 & 16,960 M^-1^ cm^-1^, Tctex = 649,850 & 31,970 M^-1^ cm^-1^, LC8 = 495,640 & 8,480 M^-1^ cm^-1^, and LC7 = 533,510 & 6,990 M^-1^ cm^-1^. All purified proteins were stored at 4°C with a protease inhibitor mixture of pepstatin A and phenylmethanesulfonyl fluoride and used within one week.

### Full Length IC Cloning and Expression

Full length CT IC (IC_FL_) was expressed in Insect Sf9 cells. The sequence of IC_FL_ was codon optimized and cloned into pFastbac1 vector by Genscript (Piscataway, NJ) with an N-terminal 6×His tag followed by a TEV protease cleavage site. IC_FL_ was expressed in Sf9 cells using the multiBAC system following previously published protocols with slight modifications (7, 57). The plasmid was transformed into EmBacY cells (Multibac) and single, white colonies were selected after 2 days and inoculated into 2xTY media supplemented with antibiotics and grown overnight. The bacteria pellet was harvested at 4000 rpm for 10 min and resuspended in 0.3 mL QIAGEN miniprep buffer P1, followed by 0.3 mL P2 buffer. After 5 min incubation, 0.4 mL P3 buffer was added and the mixture was centrifuged at 13,000 rpm for 10 min. The supernatant was added to 0.8 mL ice cold isopropanol and bacmid DNA was pelleted for 10 min at 13,000 rpm. The pellet was washed twice with 70% ethanol, resuspended in H_2_O, and stored at 4°C.

Sf9 cells were cultured in SF-900 III SFM (Thermo Fisher) at a shaking speed of 125 rpm at 27°C. 2 µg fresh bacmid DNA was transfected into 2 mL Sf9 cells at 0.9×10^6^ cells/2 mL with 8 µL Cellfection II (Thermo Fisher) following manufacturer’s protocol. Five days later, 0.5 mL of the transfected culture medium was added to a 50 mL culture of Sf9 cells (0.5×10^6^ cells/mL) for P2 infection. Four days later, cells were spun down at 3000 rpm for 15 min at 4°C and the supernatant of P2 virus was collected and stored at 4°C in the dark. Protein expression was induced by adding P2 virus to Sf9 cells (1:100 ratio, 2×10^6^ cells/mL). After four days, cells were harvested by centrifugation at 4000 rpm for 20 min at 4°C. The pellet was flash frozen in liquid nitrogen and stored at −80°C for further protein purification. Affinity purification was carried out as described above for other constructs.

### IC Structure Prediction

Sequences for IC from a range of species were obtained from the UniProt protein database (65). *Rattus norvegicus* (rat, UniProt: Q62871-3), *Drosophila melanogaster* (fruit fly, UniProt: Q24246-11), *Saccharomyces cerevisiae* (yeast, UniProt: P40960-1), *Homo sapiens* (human, UniProt: O14576-2), *Danio rerio* (zebrafish, UniProt: A1A5Y4-1), *Callorhinchus milii* (Australian ghost shark, UniProt:V9KAN3-1), *Octopus bimaculoides* (Californian two-spot octopus, UniProt:A0A0L8HM30-1), *Caenorhabditis elegans* (nematode, UniProt: O45087-1), and *Chaetomium thermophilum* (CT, thermophilic fungus, UniProt: G0SCF1-1). The first 260 amino acids of the IC from each species were scored using the Agadir algorithm, which outputs a prediction for percent helicity per residue (46,47,49–51). Agadir is not developed for prediction of long protein sequences, therefore results were compared to predictions from PSIPRED (45, 48) and in all cases agreed well.

### Circular Dichroism

CD measurements were made using a JASCO (Easton, Maryland) J-720 circular dichroism spectropolarimeter. Samples consisted of proteins at concentrations of 5-10 µM in 10 mM sodium phosphate buffer (pH 7.5). All experiments were done using a 400 µL cuvette with a path length of 0.1 cm. The data shown are the average of three scans. Thermal unfolding data were collected in increments of 5°C over a temperature range of 5 to 60°C. CD measurements were used to estimate the fractional helicity of the samples using the equation below:

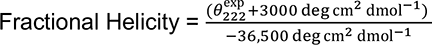

where 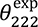 is the experimentally observed residue ellipticity (MRE) at 222 nm (52).

### Isothermal Titration Calorimetry

ITC experiments were conducted using a MicroCal VP-ITC microcalorimeter (Malvern Panalytical, United Kingdom). All experiments were performed at 25°C and with protein samples in a buffer containing 50 mM sodium phosphate (pH 7.5), 50 mM sodium chloride, and 1 mM sodium azide. Samples were degassed at 25°C prior to loading. Each experiment was started with a 2 µL injection, followed by 27 to 33 injections of 10 µL. Protein concentrations in the cell ranged from 20-40 μM and concentrations in the syringe ranged from 200-400 μM. The data were processed using Origin 7.0 (Malvern Panalytical, Malvern, UK) and fit to a single-site binding model. The recorded data are the averages of 2-3 independent experiments.

### NMR measurements and analysis

Samples for NMR were prepared in a 20 mM sodium chloride, 50 mM phosphate (pH 7.4) buffer that included 5% D_2_O, 1 mM sodium azide, 0.2 mM 2,2-dimethylsilapentane-5-sulfonic acid (DSS), and 1× cOmplete™ protease inhibitor cocktail (Roche, Basel, Switzerland). NMR spectra were collected over a temperature range of 10-40 °C using a Avance III HD 800 MHz spectrometer (Bruker Biospin, Billerica, Massachusetts) with a TCI cryoprobe and a Avance NEO 600 MHz spectrometer (Bruker Biospin) equipped with a room-temperature TXI probe. Band-selective excitation short transient (BEST) variants of TROSY-based triple resonance sequences (HNCO, HNCA, HN(CO)CA, HN(CA)CO, HNCACB, HN(CO)CACB were used for backbone assignment of IC_1-260_ at 10°C (66). Assignments of ^15^N-labeled IC_216-260_ was carried out using ^15^N-seperated TOCSY and NOESY experiments. The accessibility of IC_1-260_ amide protons to exchange with the solvent was determined by measuring peak volumes in a Fast HSQC spectrum with a 20 ms CLEANEX-PM mixing period. The IC_1-260_ and IC_216-260_ concentrations were 350-500 µM for these samples.

NMR experiments of binary complexes (IC with one binding partner) were performed by combining ^15^N-labeled IC_1-260_ with unlabeled p150_CC1B_, NudE_CC_, Tctex, LC8, or LC7 at a molar ratio of 1:1.5. For the IC_216-260_/IC_216-260_ binary complex, the ^15^N-labeled components were mixed in a 1:1 molar ratio and each was at a concentration of 350 µM. Spectra for each binary complex were collected at both 10 and 40°C. NMR experiments of the dynein subcomplex (IC_1-260_ with all light chains) were performed much in the same way. The IC_1-260_ concentration was 250 µM for these samples. NMR data were processed using TopSpin 3.6 (Bruker) and NMRPipe (67). For 3D experiments that employed non-uniform sampling the spectra were reconstructed using SCRUB (68). Peak assignment was performed using CCPN Analysis 2.5.2 (69).

### Analytical ultracentrifugation

Sedimentation velocity analytical ultracentrifugation (SV-AUC) experiments were performed using a Beckman Coulter Optima XL-A analytical ultracentrifuge, equipped with absorbance optics. For individual proteins, the concentration used was 15-30 µM. For binary complexes, IC_1-260_ or IC_FL_ were mixed with each binding partner at ratios of 1:1.5 or 1:2 (molar ratio of IC to binding partner). The buffer condition used for all SV-AUC experiments was 25 mM tris(hydroxymethyl)aminomethane hydrochloride (pH 7.4), 150 mM KCl, 5 mM tris (2-carboxyethyl) phosphine, and 1 mM sodium azide. Samples were loaded into standard, 12 mm pathlength, 2-channel sectored centerpieces and centrifuged at 42,000 rpm and 20°C. 300 scans were acquired at 280-297 nm with no interscan delay. Data were fit to a c(S) distribution using SEDFIT (70).

Sedimentation equilibrium analytical ultracentrifugation (SE-AUC) experiments were performed on the same instrument. NudE_CC_ and p150_CC1B_ were each loaded with three concentrations in the range of 15-60 μM in 6 channel centerpieces, centrifuged at three speeds (10,000, 14,000, and 18,000 rpm) and scanned at 280 nm. Samples were scanned every 3 hours until they were at equilibrium (i.e., when the final sequential scans were superimposable), which occurred after 30 to 36 hours of centrifugation. Data were acquired as averages of five measurements of absorbance at each radial position, with a nominal spacing of 0.003 cm between each position. The data from the three speeds and three concentrations were globally fit to a monomer-dimer self-association model and resulted in random residuals. Other models tested did not give adequate variances and random residuals. All experiments were done at 20°C. Data were fit using HETEROANALYSIS (71).

### SEC multiangle light scattering

SEC coupled to multiangle light scattering (SEC-MALS) was carried out using a Superdex 200 gel filtration column on an AKTA fast liquid chromatography system (Cytiva Life Sciences) coupled with a DAWN multiple-angle light scattering detector and an Optilab refractive index detector (Wyatt Technology, Santa Barbara, CA). Data for IC_1-260_ was collected for protein samples at a concentration of 200 µM protein in a buffer composed of 50 mM sodium phosphate (pH 7.5), 50 mM sodium chloride, and 1 mM sodium azide. Data for IC_FL_ was collected for protein samples at a concentration of 30 µM in a buffer composed of 25 mM tris(hydroxymethyl)aminomethane hydrochloride (pH 7.4), 150 mM KCl, 5 mM β-mercaptoethanol, and 1 mM sodium azide. Molar mass and error analysis were determined using ASTRA v9, employing a Zimm light scattering model (Wyatt Technology).

### Subcomplex reconstitution

Post purification of individual protein, IC_1-260_ was combined with dynein light chains (Tctex, LC8, and LC7). To the assembled subcomplex, the non-dynein proteins p150_CC1B_ or NudE_CC_ were added in a 1:1.5 molar ratio prior to SEC using a Superdex 200 column (Cytiva Life Sciences). Subcomplex reconstitution was verified by SDS-PAGE of SEC fractions, as each protein clearly resolves. When estimating concentrations for complexes purified by SEC, the absorbance at 280 nm was used with the assumption that the majority of formed complex in solution followed the expected stoichiometry of 1:1 (IC monomer: partner monomers) and that very little excess of free protein would be present.

## Acknowledgements

We acknowledge support from the National Science Foundation (Award 1617019 for E.J.B. and Award 2003557 for N.M.L.). The Oregon State University NMR Facility is funded in part by the National Institutes of Health (HEI Grant 1S10OD018518) and by the M. J. Murdock Charitable Trust (Grant 2014162). The Lewis & Clark College Bruker Avance NEO 600 NMR spectrometer was purchased with support from the National Science Foundation (Award 1917696) and the M. J. Murdock Charitable Trust (Grant 201811283).

---

**Figure S1:**
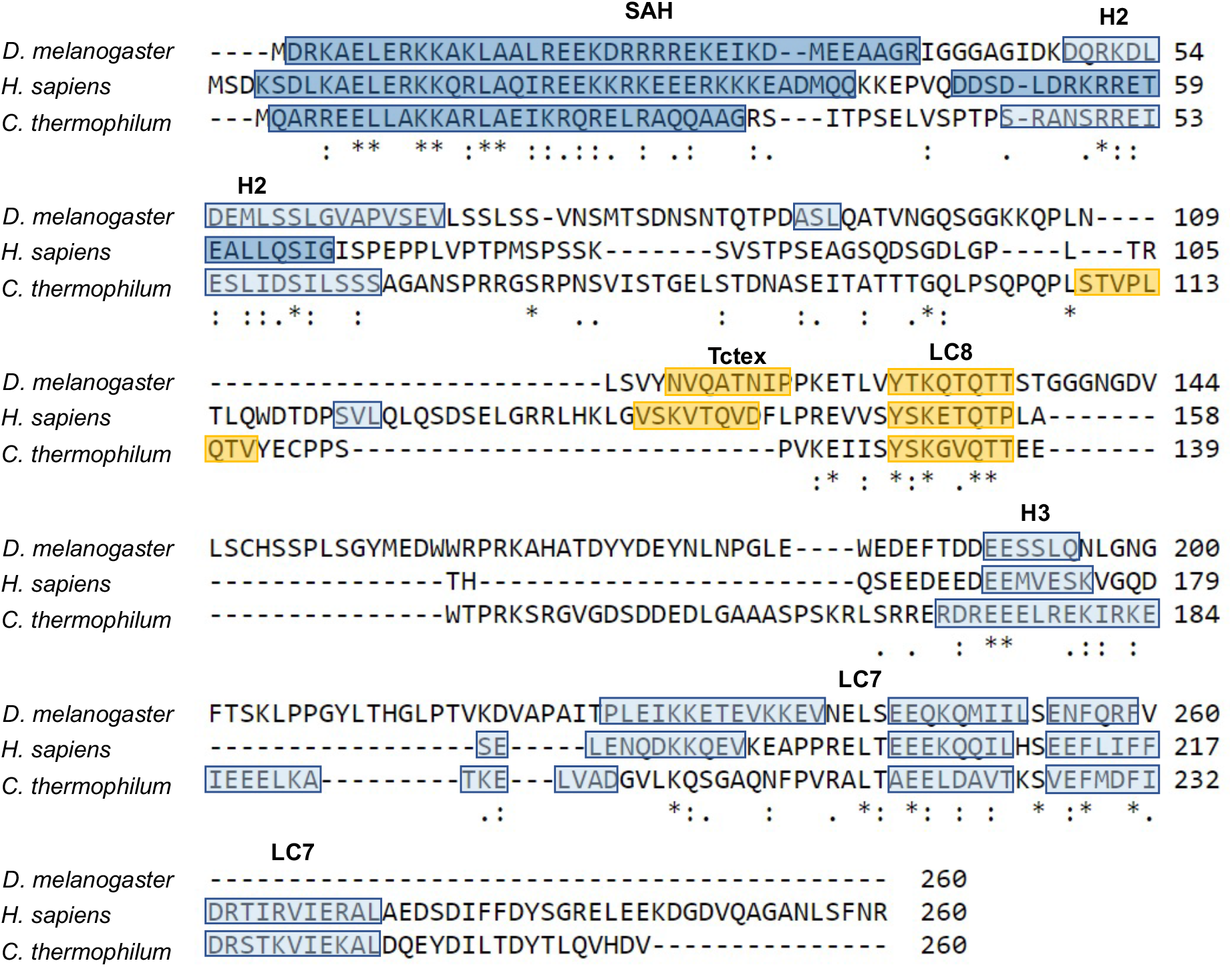
IC sequence alignment. Alignment of the first 260 residues of IC from Drosophila, human, and CT using the MAFFT alignment program (72). Identical (asterisk), strongly similar (colon), and weakly similar (period) residues are shown at the bottom of each alignment. Known or predicted α-helical secondary structure (SAH, H2, H3, and the LC7 binding domain) is highlighted in blue, with the darker shade of blue indicating stronger prediction. Known or predicted binding sites for Tctex and LC8 are highlighted in yellow (73).

**Figure S2:**
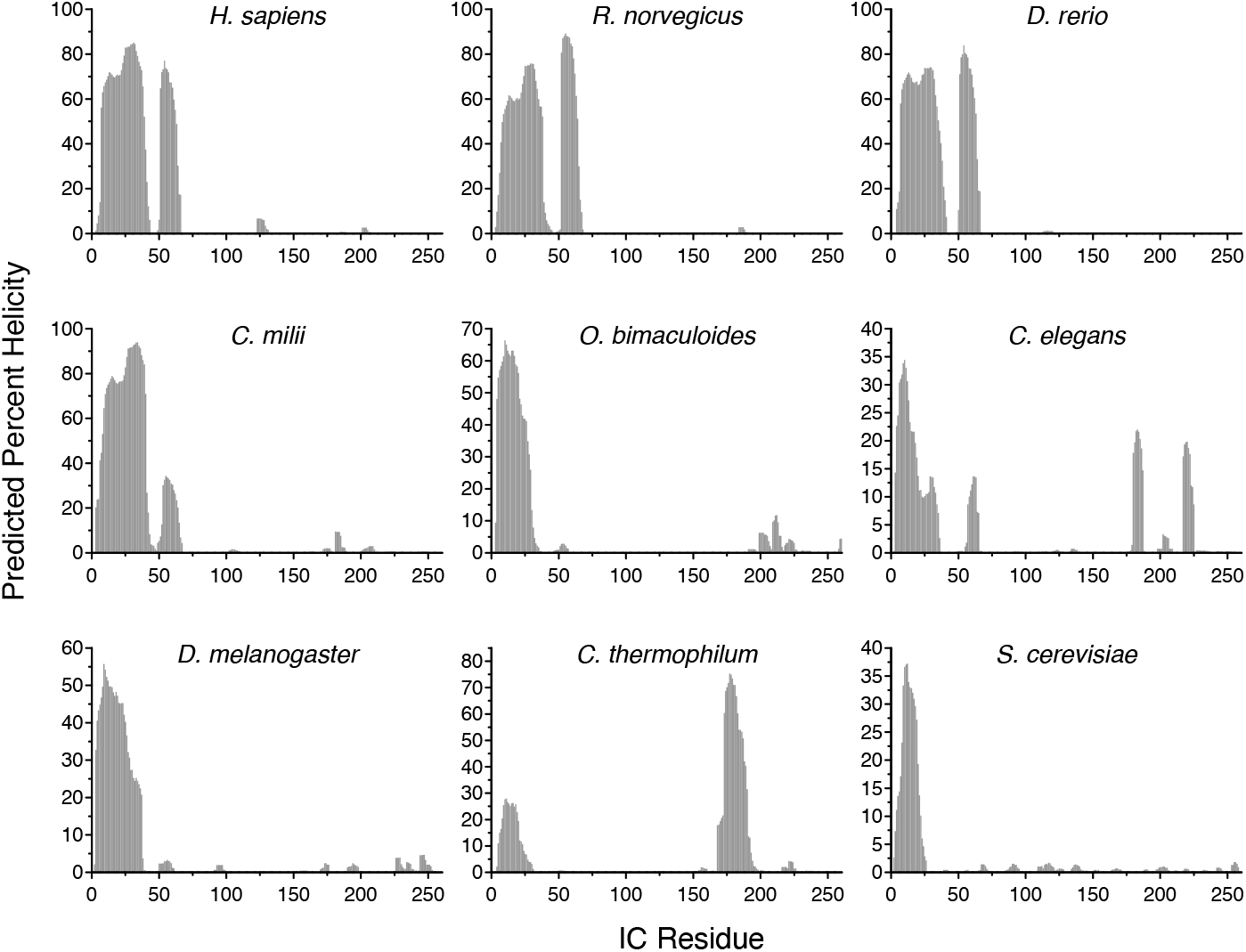
Predicted percent Helicity in IC across species. Residue-level percent helicity predictions generated using the Agadir algorithm for the first 260 amino acids of IC from *H. sapiens* (human), *R. norvegicus* (rat), *D. rerio* (zebrafish), *C. milii* (Australian ghostshark), *O. bimaculoides* (Californian two-spot octopus), *C. elegans* (nematode), *D. melanogaster* (fruit fly), *C. thermophilum* (CT, thermophilic fungus), and *S. cerevisiae* (yeast) (46,47,49–51). **Figure S2-source data 11: Source file for helix prediction of IC across species.** This Excel workbook contains the data plotted for the helix predictions shown in Figure S2. Additional information regarding data collection can be found in the corresponding methods section. Data were plotted using gnuplot.

**Figure S3:**
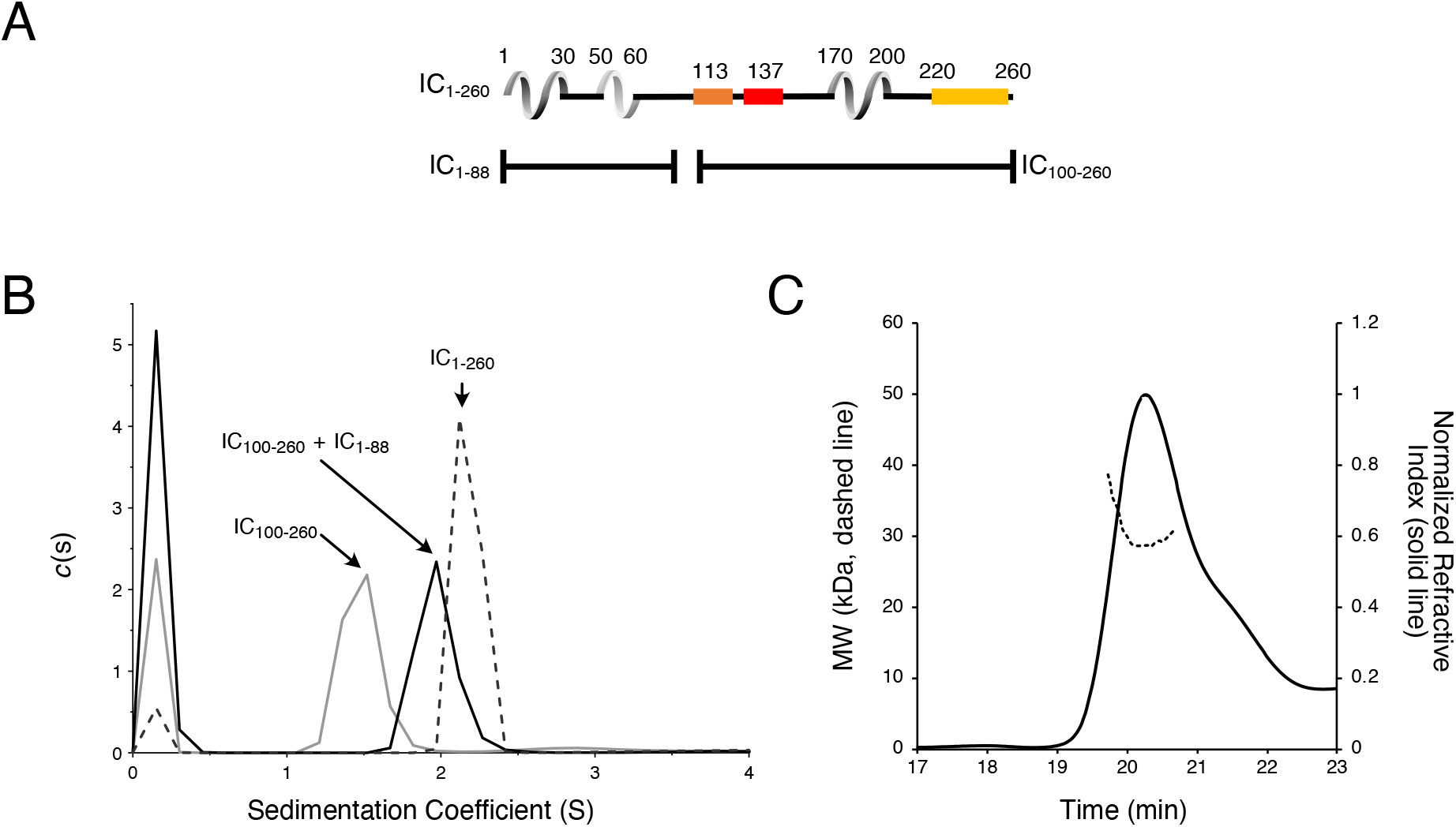
IC1_-88_ and IC_100-260_ binding by SV-AUC. (A) Domain architecture diagrams of IC_1-260_, IC_1-88_, and IC_100-260_. (B) SV-AUC experiments of IC_100-260_ (grey), IC_100-260_ mixed with IC_1-88_ at a 1:2 molar ratio (black), and IC_1-260_ (grey dashes). (C) The estimated mass of IC_100-260_/IC_1-88_ complex from MALS is 30.3 kDa, which indicates a 1:1 binding stoichiometry. **Figure S3-source data 12: Source file for SV-AUC and SEC-MALS of binding interactions between IC_1-88_ and IC_100-260_.** This Excel workbook contains the data plotted for the SV-AUC and SEC-MALS data shown in Figure S3 with different sheet corresponding to different panels within the figure. Additional information regarding data collection can be found in the corresponding methods section. Data were plotted using gnuplot.

**Figure S4:**
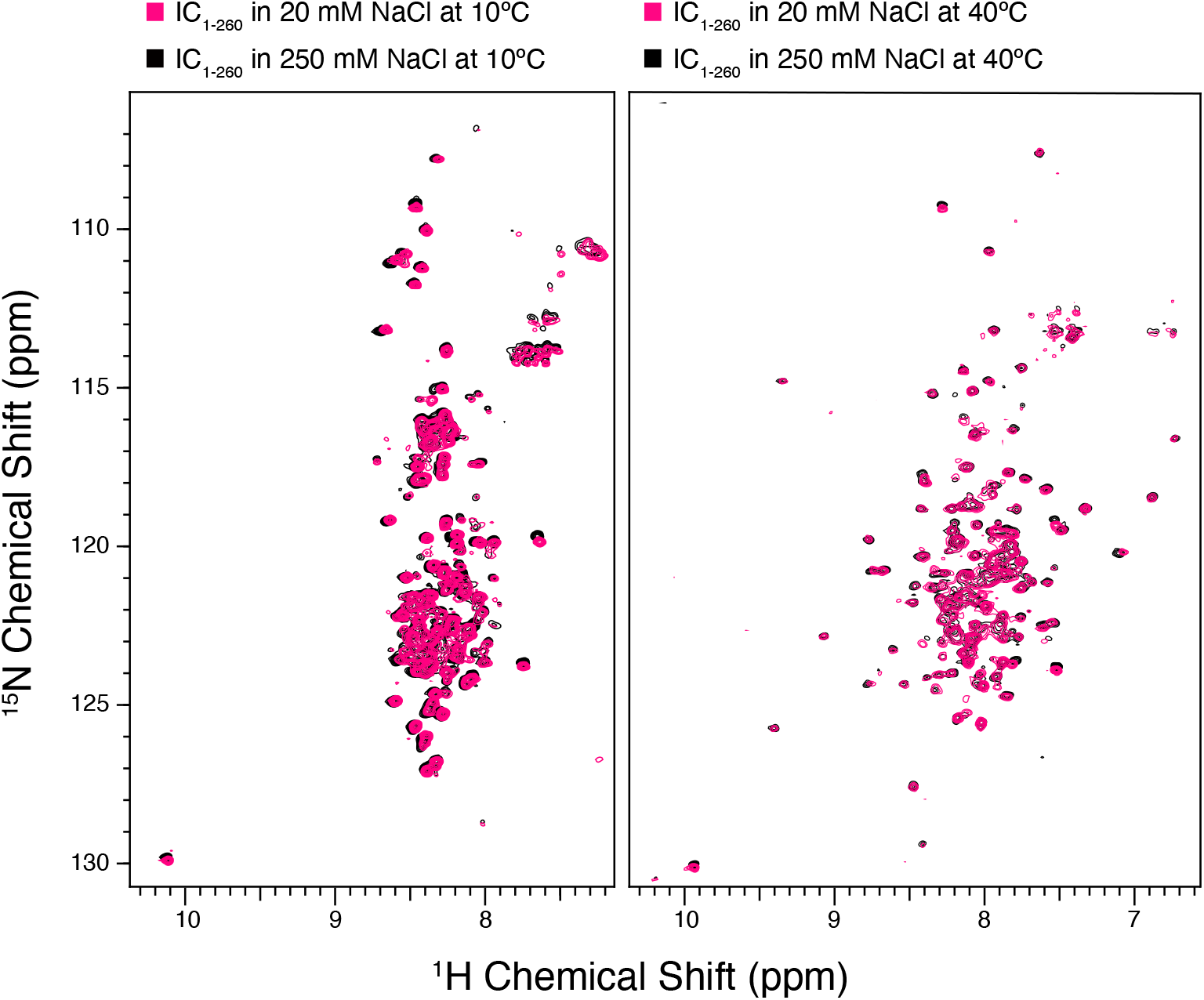
NMR spectra of IC are unaffected by salt concentration. ^1^H-^15^N TROSY spectra of IC_1-260_ in 20 mM NaCl (black) overlaid with IC_1-260_ in 250 mM NaCl (pink) at both 10°C (left) and at 40°C (right).

